# A complete electron microscopy volume of the brain of adult *Drosophila melanogaster*

**DOI:** 10.1101/140905

**Authors:** Zhihao Zheng, J. Scott Lauritzen, Eric Perlman, Camenzind G. Robinson, Matthew Nichols, Daniel Milkie, Omar Torrens, John Price, Corey B. Fisher, Nadiya Sharifi, Steven A. Calle-Schuler, Lucia Kmecova, Iqbal J. Ali, Bill Karsh, Eric T. Trautman, John Bogovic, Philipp Hanslovsky, Gregory S. X. E. Jefferis, Michael Kazhdan, Khaled Khairy, Stephan Saalfeld, Richard D. Fetter, Davi D. Bock

**Affiliations:** Janelia Research Campus, Howard Hughes Medical Institute, Ashburn, VA 20147; Coleman Technologies, Inc., Newtown Square, PA 19073; Hudson Price Designs, LLC, Hingham, MA 02043; Division of Neurobiology, MRC Laboratory of Molecular Biology, Cambridge, CB2 0QH, UK; Department of Computer Science, Johns Hopkins University, Baltimore, MD 21218

**Keywords:** Electron microscopy, connectomics, neural circuits, *Drosophila melanogaster*, mushroom body, olfaction, image stitching

## Abstract

*Drosophila melanogaster* has a rich repertoire of innate and learned behaviors. Its 100,000–neuron brain is a large but tractable target for comprehensive neural circuit mapping. Only electron microscopy (EM) enables complete, unbiased mapping of synaptic connectivity; however, the fly brain is too large for conventional EM. We developed a custom high-throughput EM platform and imaged the entire brain of an adult female fly. We validated the dataset by tracing brain-spanning circuitry involving the mushroom body (MB), intensively studied for its role in learning. Here we describe the complete set of olfactory inputs to the MB; find a new cell type providing driving input to Kenyon cells (the intrinsic MB neurons); identify neurons postsynaptic to Kenyon cell dendrites; and find that axonal arbors providing input to the MB calyx are more tightly clustered than previously indicated by light-level data. This freely available EM dataset will significantly accelerate *Drosophila* neuroscience.

**HIGHLIGHTS:**

- A complete adult fruit fly brain was imaged, using electron microscopy (EM)
- The EM volume enables brain-spanning mapping of neuronal circuits at the synaptic level
- Olfactory projection neurons cluster more tightly in mushroom body calyx than expected from light-level data
- The primary postsynaptic targets of Kenyon cells (KCs) in the MB are other KCs, as well as the anterior paired lateral (APL) neuron
- A newly discovered cell type, MB-CP2, integrates input from several sensory modalities and provides microglomerular input to KCs in MB calyx
- A software pipeline was created in which EM-traced skeletons can be searched for within existing large-scale light microscopy (LM) databases of neuronal morphology, facilitating cell type identification and discovery of relevant genetic driver lines

## INTRODUCTION

Neural circuits are in large part made of neurons and the synapses connecting them. Maps of connectivity inform and constrain all models of how neuronal circuits transform information and subserve behavior (Braitenberg and Schüz, 1998; Marr, 1969; Sterling and Laughlin, 2015). Historically, anatomical maps of neuronal connectivity were inferred from light microscopy (LM) images of sparsely labeled neurons (Shepherd, 2016). Updated forms of this approach remain important to this day (*e.g.* Wertz et al., 2015; Wolff et al., 2015), as do electrophysiological measurements of connectivity between small groups of neurons (Ko et al., 2011; Perin et al., 2011; Song et al., 2005). However, for a given volume of brain tissue, these methods lack the resolution to map all synapses between all neurons, which may result in an undersampled description of neuronal network topology (Helmstaedter et al., 2008).

Electron microscopy (EM) is the only method capable of simultaneously resolving all neuronal processes and synapses in a given volume of brain tissue – a requirement if one wishes to make complete maps of neuronal connectivity at the synapse level (or ‘connectomes’; Lichtman and Sanes, 2008). However, generating EM volumes of any appreciable size is technically challenging (Briggman and Bock, 2012; Harris et al., 2006; Helmstaedter, 2013). Nanometer-scale image voxels must be acquired over a spatial extent sufficient to encapsulate circuits of interest, typically tens to hundreds of microns at a minimum. Volume EM for connectomics has therefore traditionally been limited to exceedingly small organisms, such as the nematode (White et al., 1986) and the larval ascidian (Ryan et al., 2016), or to small subvolumes from (for example) the fly optic medulla (Takemura et al., 2008), cat thalamus (Hamos et al., 1987), and macaque visual cortex (McGuire et al., 1991).

Recent technical advances have enabled increased acquisition speed and automation of the imaging pipeline, producing larger EM volumes than were previously attainable (reviewed in Briggman and Bock, 2012; see also Eberle et al., 2015; Kuwajima et al., 2013; Xu et al., 2017). Circuit diagrams mapped in these larger EM volumes have yielded new insights into (for example) the network architecture of the larval fly (Ohyama et al., 2015), the optic medulla of the adult fly (Takemura et al., 2017b), the zebrafish olfactory bulb (Wanner et al., 2016), and the mammalian retina (Briggman et al., 2011; Lauritzen et al., 2016), thalamus (Morgan et al., 2016), and neocortex (Bock et al., 2011; Kasthuri et al., 2015; Lee et al., 2016). Large EM volumes have also revealed surprising new findings in cellular neuroanatomy, such as the differential distribution of myelin on axons depending on neuronal subtype (Micheva et al., 2016; Tomassy et al., 2014). However, imaging infrastructure for volume EM continues to limit the scale of connectomics investigations.

Here we report next-generation hardware and software for high throughput acquisition and processing of EM data sets. We apply this infrastructure to image the entire brain of a female adult fruit (aka vinegar) fly, *Drosophila melanogaster* (Figure 1A). At approximately 8 × 10^7^ μm^3^, this volume is nearly two orders of magnitude larger than the next-largest complete brain imaged at sufficient resolution to trace synaptic connectivity, that of the first instar *Drosophila* larva (Ohyama et al., 2015).

**Figure 1.**
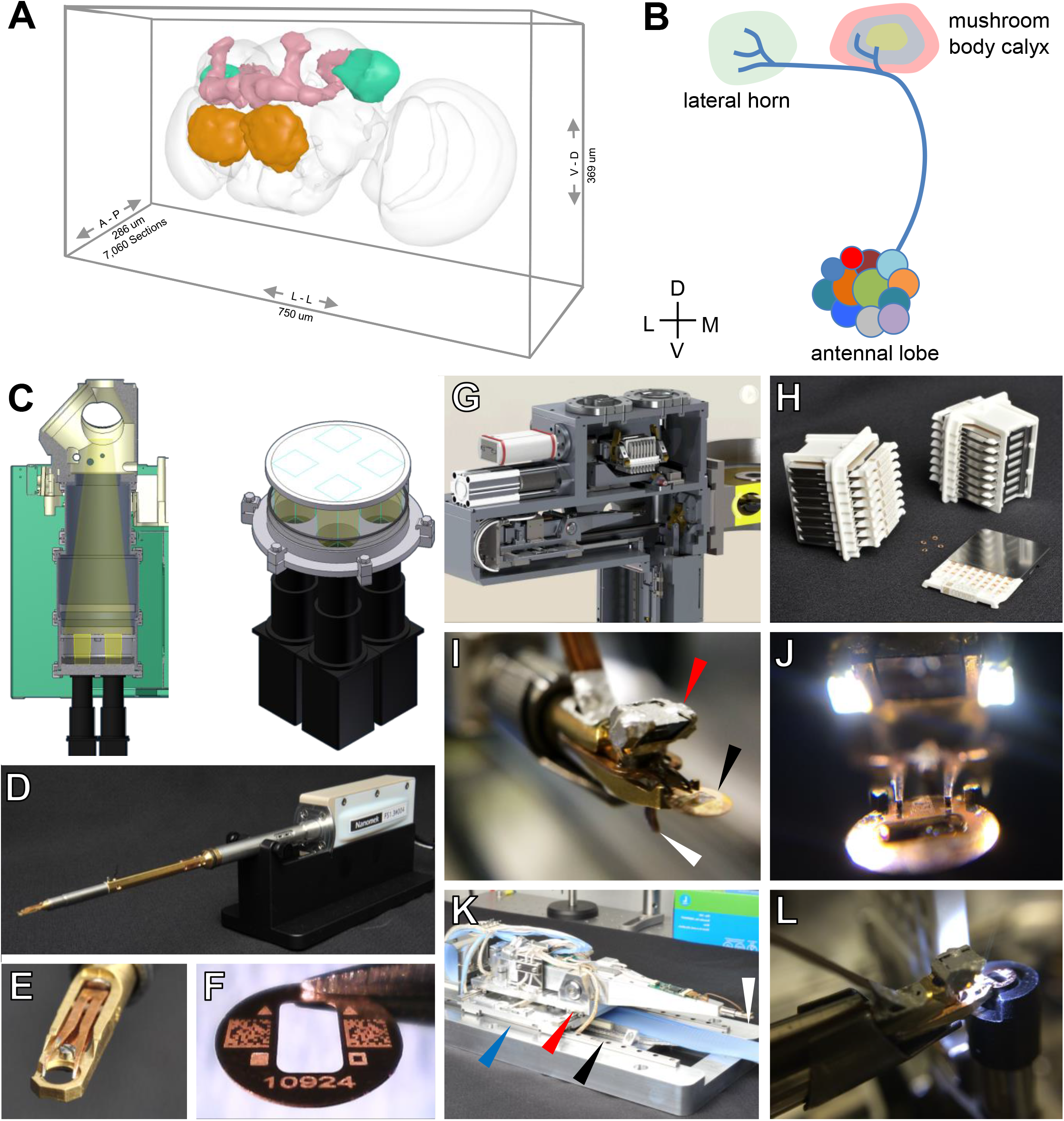
Target Volume and EM Acquisition Infrastructure. See also Figure S1, Figure S2, Figure S3, Movie S1, Movie S2, Movie S3. (A) Oblique view of a surface model of the *Drosophila* brain (gray mesh) with specific neuropils highlighted: antennal lobe (orange); mushroom body (pink); lateral horn (turquoise). (B) Schematic of olfactory pathway. Projection neurons (PNs) originate from antennal lobe and their axons pass through the MB calyx, forming *en passant* synapses with MB output neurons (MBONs), before terminating in the LH. (C) Left, schematic of TEMCA2 vacuum extension, scintillator, and camera array. Right, camera field of views (FOVs) diagram, indicating the non-overlapping FOV of each camera on scintillator. (D) FEI CompuStage-compatible single-axis Fast Stage. (E) Fast Stage grid holder. (F) Custom-etched 2×1 mm slot grid with fiducial marks, 2-D barcodes, and unique serial identifier. (G) Cassette, magazines, and four-axis stage inside the Autoloader vacuum. (H) Autoloader cassettes and magazines. (I) Grid holder and end-effector of Autoloader grid positioning system (GPS). Arrows: prism and LED assembly (red); sample grid (black); lever of the grip assembly which actuates grid release when retracted against its stop on the stage housing (white). (J) Autoloader end-effector (K) Four-axis Autoloader stage. Arrows: grid positioning system (GPS) chamber (blue); view port (red); cassette shuttle chamber (black); end effector and miniature camera (white). (L) Rotational aligner integrated into the Autoloader cassette shuttle.

*D. melanogaster* is an important model organism for neurobiology research, owing to its rich repertoire of innate and learned behavior (Hampel et al., 2015; Heisenberg and Wolf, 1984; Hoopfer, 2016; Kim et al., 2017; Ofstad et al., 2011; Owald and Waddell, 2015; Pavlou and Goodwin, 2013; von Reyn et al., 2014), electrophysiological accessibility (*e.g.* Hige et al., 2015; Wilson et al., 2004), relatively small size (Figure 1A), and the stereotypy of and genetic access to most of the ~100,000 neurons in its brain (Aso et al., 2014; Chiang et al., 2011; Jenett et al., 2012; Kvon et al., 2014; Milyaev et al., 2012; Pfeiffer et al., 2010). In the fly brain, each morphological cell type usually consists of one to a few neurons per hemisphere, with stereotyped neuronal arbors reproducible across individuals with a precision of ~10 microns (Costa et al., 2016; Jefferis et al., 2007; Lin et al., 2007). Thousands of genetic driver lines for specific subsets of cell types (Jenett et al., 2012; Kvon et al., 2014), or even single cell types (Aso et al., 2014; Grabe et al., 2015; Wolff et al., 2015), enable *in vivo* manipulation of neuronal physiology and the construction of searchable databases of neuronal morphology (Chiang et al., 2011; Costa et al., 2016; Milyaev et al., 2012).

We leveraged the stereotypy of fly neuronal morphology to validate that the EM volume was suitable for tracing brain-spanning neuronal circuitry. We focused on the olfactory projection neurons (PNs), which are thoroughly described at the light microscopy (LM) level (Jefferis et al., 2007; Lin et al., 2007; Tanaka et al., 2004) (Figure 1B, Figure S1). On each side of the brain, the dendrites of ~150 PNs innervate ~50 glomeruli of the antennal lobe (AL). Each glomerulus is morphologically identifiable (Couto et al., 2005; Grabe et al., 2015; Stocker et al., 1990) and receives input from a stereotyped set of olfactory receptor neurons (ORNs), resulting in reproducible PN odorant tunings across animals (Wilson, 2013; Wilson et al., 2004). PN axons project from the AL to the lateral horn (LH), which is thought to subserve stereotyped behavioral responses to odorants (Heimbeck et al., 2001; Jefferis et al., 2007; Ruta et al., 2010). Along the way to the LH, most PNs send collaterals into the calyx of the mushroom body (MB), a locus of learning, recall, and synaptic plasticity (Davis, 2011; Heisenberg, 2003; Owald and Waddell, 2015). Most PN types project to the MB calyx via the medial antennal lobe tract (mALT), but several travel in secondary tracts, and a few bypass calyx entirely and project only to LH (Frank et al., 2015; Stocker et al., 1990; Tanaka et al., 2012).

To explore whether the EM volume could be used to make new discoveries as well as verify existing knowledge, we examined a subset of the circuitry downstream to PNs in the MB calyx. The *Drosophila* MB has ~2,000 intrinsic neurons on each side of the brain called Kenyon cells (KCs). Each KC projects a highly variable dendritic arbor into the calyx, which terminates in elaborations known as claws (Figure S1). Claws from many KCs converge to wrap individual PN boutons in a characteristic structure called the microglomerulus (Yasuyama et al., 2002), and each KC receives input from multiple PNs of diverse types (Caron et al., 2013; Gruntman and Turner, 2013). KCs sample this input in what is thought to be a random fashion (Caron et al., 2013), although some biases have been noted (Gruntman and Turner, 2013). In order to fire action potentials, KCs require a threshold number of input PNs to be coactive (Gruntman and Turner, 2013); the firing pattern of KCs is therefore thought to be a combinatorial and sparse representation of olfactory stimuli. The dendrites of KCs also receive inhibitory and modulatory synapses from a variety of other cell types within the calyx, and have presynaptic release sites, which target unknown cell types (Butcher et al., 2012; Christiansen et al., 2011). KC axons project from the calyx to the MB lobes, where they synapse onto MB output neurons (MBONs). KC-MBON synapses are modulated by rewarding or punishing signals from dopaminergic afferent neurons (DANs; Aso et al., 2012; Burke et al., 2012; Liu et al., 2012); this plasticity underlies olfactory memory formation (Hige et al., 2015; Owald et al., 2015; Sejourne et al., 2011).

In the current work, we surveyed all microglomeruli in the main MB calyx and traced their bouton inputs sufficiently to identify the originating cell’s type, resulting in a description of the complete set of olfactory inputs to the MB. Although most MB input originated from olfactory PNs, we discovered a previously unknown cell type providing bouton input to KC claws. To map unknown connectivity within the calyx, we also identified the cell types of the KC postsynaptic targets. Finally, we found more clustering of PN axonal afferents within the MB calyx than was predicted from light microscopy (LM) data, which may bias the sampling of olfactory input from PNs by KCs. These findings demonstrate the utility of this whole-brain dataset for mapping both known and new neural circuit connections.

## RESULTS

### TEMCA2: a second-generation transmission EM camera array

Current volume EM methods generally trade off between convenient sample handling, high image resolution, and rapid image acquisition (Briggman and Bock, 2012). Transmission EM (TEM) camera arrays (TEMCAs) offer high signal-to-noise and high throughput imaging of serial thin sections (Bock et al., 2011; Lee et al., 2016). Post-section staining increases sample contrast over alternative methods that rely on *en bloc* staining, and features of interest may be re-imaged at higher magnifications. However, lossless serial sectioning and imaging of thousands of sections is a technically challenging, manual process; the image data are anisotropic (i.e., each voxel is narrower than it is tall, typically 4 × 4 × 45 nm), which is inconvenient for processing by automated segmentation pipelines; and large sample areas represented by mosaics of thousands of overlapping individual camera images necessitate a sophisticated and scalable stitching pipeline (Saalfeld et al., 2012; Wetzel et al., 2016).

Despite these challenges, the potential for gains in throughput persuaded us to develop a second-generation system (TEMCA2) prior to undertaking a *Drosophila* whole-brain imaging effort (Figure 1; Figure S2). To achieve high frame rates in TEM, electron dose is simply increased until the sensors are saturated in the desired image frame integration period. This option is generally not available in scanning EM (SEM)-based approaches, since coulomb repulsion between electrons limits the maximum current per beam (Denk and Horstmann, 2004). We constructed a 2 × 2 array of high frame-rate sCMOS-based scientific cameras and coupled them with optical lenses custom-designed for imaging TEM scintillators (Figure 1C). At this higher electron dose, images are acquired in 4 frames of 35 ms exposure each. Standard TEM sample holders and goniometer stages take several seconds to step and settle, which is not fast enough for this imaging scheme. Therefore, we built a high speed, piezo-driven single axis Fast Stage (Figure 1D; Figure S2B; Movie S1), with a sample holder designed to accept standard-diameter (3 mm) TEM sample grids (Figure 1E-F). The Fast Stage has the same shape as a standard sample holder, so that the TEM’s standard multi-axis stage can provide motion in other axes. Step-and-settle with the Fast Stage typically completed in 30-50 ms (Figure S2C-D). On-line analysis of sample drift between subsequent frames was used to determine whether stability was sufficient to acquire high-quality images, and frames were translated before summation to correct for small (16 nm or less) drift between frames. Net imaging throughput using the TEMCA2 system is ~50 MPix/s, roughly six times faster than the first-generation TEMCA (Bock et al., 2011). For the whole-brain imaging effort, we constructed two TEMCA2 systems, yielding an order of magnitude increase relative to previously available EM imaging throughput.

### Autoloader: a hands-free robot for automatic and reliable TEM imaging

To allow unattended multi-day imaging, reduce risk to the samples, and decrease the overhead of sample exchange (10 minutes out of every 30 in typical TEMCA2 operation), we built a robotic system (Autoloader) capable of autonomous sample exchange and imaging of the TEM grids (Figure 1G; Figure S2E-F; Movies S2-3). Although automatic sample exchange systems for TEMs have been built (Lefman et al., 2007; Potter et al., 2004), their capacity and reliability were insufficient for the whole-brain imaging effort described here. The Autoloader mounts to an accessory port on the TEM, has its own vacuum system, and completely replaces the off-the-shelf stage system. To better support automatic sample handling, we made custom 100 μm-thick beryllium-copper sample support grids, each etched with unique ID numbers and spatial fiducial marks to guide machine vision-based pick-and-place software for grid exchange (Figure 1F). Each support grid is stored in a 64-pocket cassette, and each cassette is stored in an 8-cassette magazine (Figure 1H). The Autoloader grid positioning system (GPS; Figure 1I-K) provides high-speed multi-axis grid positioning. A pre-aligner is available for optimizing sample orientation (Figure 1L; Movies S2-3). Automatic grid exchange is accomplished in about 5 minutes without breaking vacuum.

### Application of EM infrastructure to image a complete adult fly brain

For a given electron dose, a higher contrast sample scatters more electrons, resulting in a higher quality image (Denk and Horstmann, 2004). We therefore optimized fixation and embedding procedures for high membrane contrast, while preserving high quality ultrastructure. A series of 7,060 sections, encompassing the entire brain, was prepared manually (Figure S3). Nearly all (99.75%) targeted serial section data were successfully acquired. Ten sections were lost prior to imaging, and regions of some sections with debris or cracks in the support film were excluded from imaging. Medium- and large-diameter neurites can still be readily traced through the missing data, with minimal anticipated impact on traced networks (Schneider-Mizell et al., 2016). The resulting EM dataset comprises ~21 million images occupying ~106 TB on disk.

The data were acquired over a period of ~16 months. Eighty-three percent of imaged sections were acquired with a TEMCA2 system (4.3 million Fast Stage moves), while the Autoloader was still in development, and 17% of imaged sections were acquired by the Autoloader (3.5 million GPS moves; ~6,800 machine vision-guided steps to pick, pre-align, and re-stow each grid). Eighty-two percent of Autoloader grid exchanges were successful; 14% were automatically halted and the grids re-stowed, usually due to variations in the manual placement of grids in the Autoloader cassettes or inhomogeneities in the support film; and 4% required manual control of the Autoloader for re-stowing. Re-stowed grids were removed from the Autoloader and imaged manually with a Fast Stage on a TEMCA2. No sections were lost or damaged due to Autoloader or Fast Stage malfunction.

The quality of acquired image data was high (Figure 2; Movie S4). Whether a given EM volume has sufficient resolution to reliably detect synapses and trace fine neuronal processes can currently only be evaluated empirically (see below). In general, however, image resolution increases not only with decreasing voxel size, but also with increasing image signal-to-noise (S/N). We found that the S/N of images in this dataset equals or exceeds that of other publicly available datasets (Figure 2G; Figure S4). Furthermore, it is straightforward to re-image targeted regions of interest in the full adult brain volume at higher magnification (Figure S5A-B).

**Figure 2.**
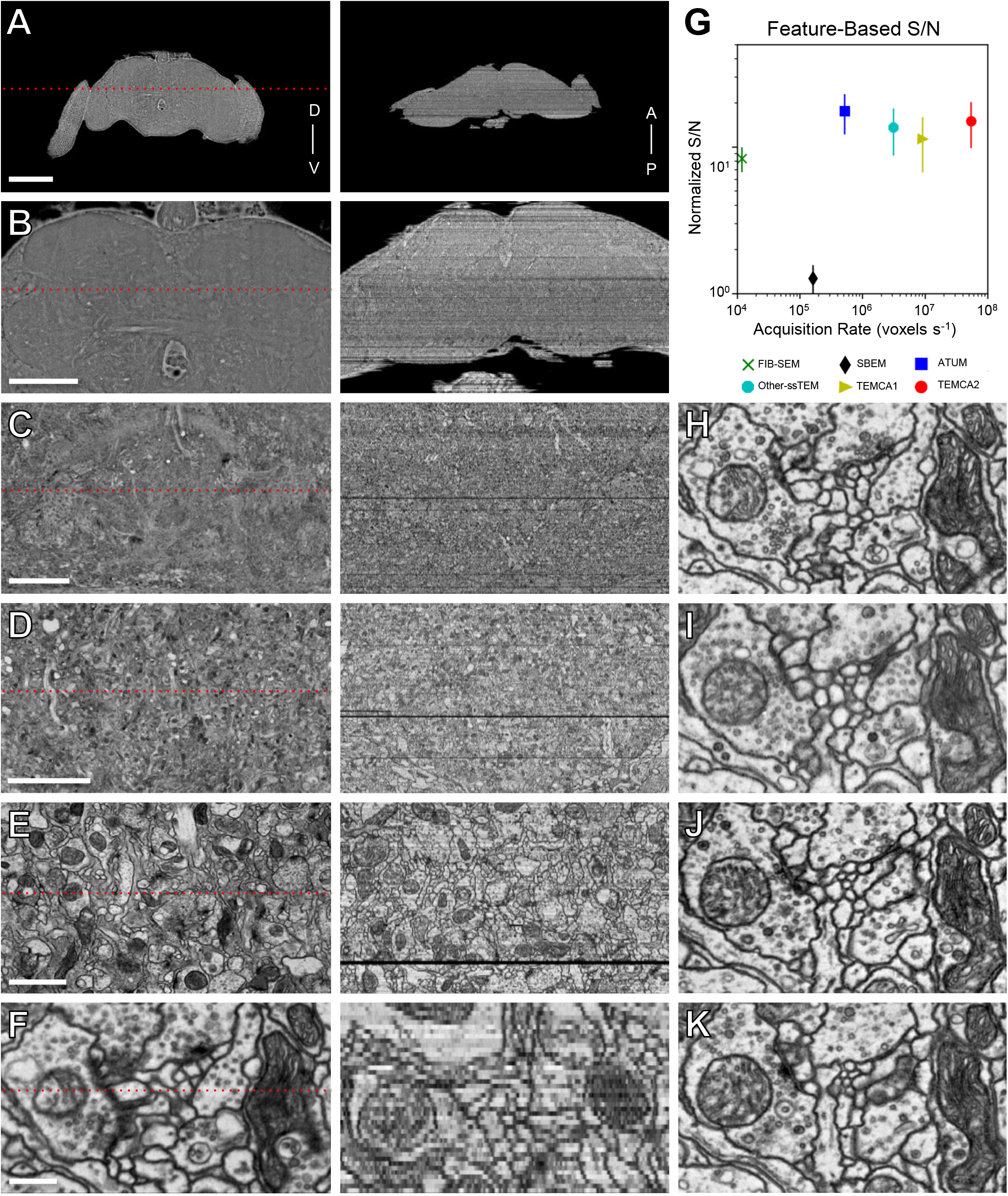
Reconstructed Image Volume. See also Figure S4, Figure S5, Figure S6. (A-F) Renderings of brain-spanning EM in the sectioning plane (x-y axes) at successive zoom levels. All panels rendered using the ELM viewer (Methods), which averages several adjacent sections to improve contrast at low magnifications. Red dotted lines in left column indicate orthogonal (y-z axes) section plane through the brain volume, rendered in right column. “D-V” and “A-P” indicate the dorso-ventral and anterior-posterior axes, respectively. (G) Image S/N versus acquisition speed for the current dataset and several publicly downloadable volume EM data sets acquired via different techniques (Table S3). Acquisition speed is in logarithmic scale. We assume all methods are shot-noise limited; for comparison purposes signal-to-noise values have therefore been normalized to the TEMCA2 voxel size (4x4x40 nm) by the square root of the source data’s voxel size (Methods). (H-K) Serial thin sections succeeding the one in F. Fine processes can be followed across serial sections and section-to-section image registration is accurate enough to provide a consistent field of view. Scale bars: 200 μm in (A), 100 μm in (B); 25 μm in (C); 10 μm in (D); 2 μm in (E); 0.4 μm in (F, H-K).

### Volume reconstruction and validation of tracing by NBLAST-based geometry matching

We developed cluster-backed software to stitch images from a single thin section into a coherent mosaic, and then to register stitched mosaics across thousands of serial sections into an aligned volume (Figure 2A-G), a process known as ‘volume reconstruction’. Calibration mosaics were used to correct lens distortions (Kaynig et al., 2010), and a scalable and linear solver was developed to stitch all section mosaics independently. During alignment of the volume, approximately 250 sections were found to be misordered. These misordered sections were automatically detected and re-ordered over several iterations of coarse and fine series alignment (Hanslovsky et al., 2017). With this software infrastructure, traced neurons can be projected across successive volume reconstructions, allowing tracing work to begin before imaging of the whole brain was complete. Furthermore, high- and low-dose imaging of robust and fragile areas of a section, respectively, could be stitched together seamlessly (Figure S5 C-E). Intra-mosaic variations in image tile intensity, created by variations in section thickness, electron beam etching, or deposition of contaminants from post-staining or section pickup, were corrected (Figure S5 F-I) using a scalable implementation of an existing algorithm (Kazhdan et al., 2010).

To test the reproducibility of tracing in the whole-brain EM dataset, three independent teams, each comprising two tracers, targeted the same KC for anatomical reconstruction (Figure S6). In the fly brain, microtubule-free neurites (‘twigs’) as fine as 40 nm in diameter tend to extend for short distances before joining larger, microtubule-containing ‘backbone’ neurites (Schneider-Mizell et al., 2016). KC claws are good examples of ‘twigs’, whereas their dendritic trunks and the axonal neurite leaving the calyx are larger-diameter ‘backbones’. The neuronal arbors and associated synapses reconstructed by each team were essentially identical for both twigs and backbones. PN to KC claw inputs with high synapse counts were detected in all three reconstructions (Figure S6C). Consistent with a tracing approach biased toward false negatives rather than false positives (Methods), one low-synapse-count input was missed by one of the tracing teams (Figure S6, red asterisks). These independent reconstructions demonstrate that the EM data support tracing of neuronal connectivity, even in challenging neuropil such as the microglomeruli of the MB calyx.

The stereotypy of the fly brain allows identification and comparison of fluorescently labeled neurons across individuals, by warping brains imaged at the light level to a standard template brain (Chiang et al., 2011; Costa et al., 2016; Manton et al., 2014; Milyaev et al., 2012). We developed tools to register LM datasets to the EM-imaged brain (Methods), allowing precise overlay of LM onto EM data across multiple brains (Figure 3A-D). This approach can also be used to analyze EM-traced neurons within existing frameworks for fly neuroanatomy. For example, the geometric search algorithm, NBLAST (Costa et al., 2016), can be used to search for an EM-traced PN skeleton thought to arise from the AL glomerulus VM2 (Figure 3E-G) in the FlyCircuit single neuron collection (Chiang et al., 2011). The VM2 PN is the top hit arising from this query (Figure 3G), with an NBLAST mean score of 0.638. Remarkably, this NBLAST score is within the range of top scores for the 6 LM-imaged VM2 neurons in the FlyCircuit database when compared with one another (0.635-0.706), consistent with the high qualitative similarity of the EM-traced and LM-imaged PNs (Figure 3G).

**Figure 3.**
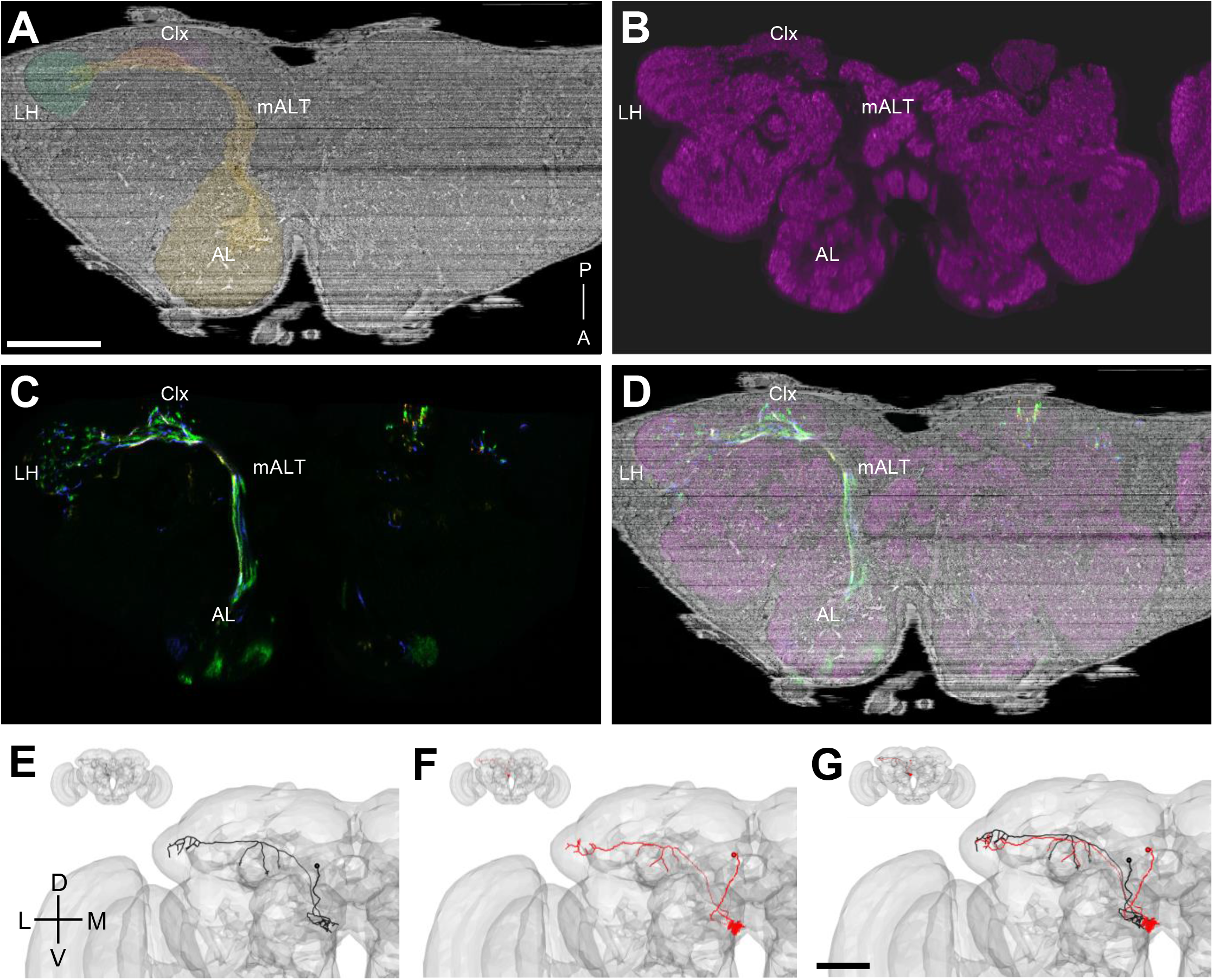
Validation of Tracing by EM-LM Registration and NBLAST-based Geometry Matching. (A-D) ELM can be used to define a three-dimensional warp field between the EM data set and a light-level template brain such that EM-imaged and LM-imaged brains are in a common template space. Same oblique cut plane shown in A-D. (A) Oblique cut plane through the EM volume contains the AL and mALT (orange) that project from AL to MB calyx (red), and LH (green). (B) The LM template brain immunofluorescently labeled with synapse-specific nc82 (magenta). The mALT is devoid of synapse-labeling. (C) LM data of a subset of PNs labeled with random fluorophore combinations using FLP-out technique. (D) Overlay of A-C. All LM datasets that have been aligned to the template brain can be projected onto the EM dataset. (E) An EM-traced putative VM2 PN (black skeleton), projected to a template brain (gray surface mesh) using the inverse of the transformation previously defined to align the template brain to the EM dataset in B. (F) Top hit resulting from an NBLAST search of the FlyCircuit database using the EM-traced PN (red) as a query structure. The annotated class in the VFB database is VM2. (G) Overlay of the EM and LM PNs shows great structural similarity. Scale bars: ~100 μm in (A-D), ~50 μm in (E-G).

### EM-based reconstruction of complete olfactory input to the MB calyx reveals tight clustering of homotypic PN arbors

To systematically compare EM-based PN reconstructions with available LM data, we identified all PN to KC microglomeruli in the main MB calyx on the right side of the fly’s brain, and traced the originating PNs sufficiently to identify their subtype (Figure 4). We classified olfactory PNs known to arise from a single glomerulus in AL based on assessment of each PN’s dendritic distribution in AL (Figure 4B) and its axonal arbor in LH. We found that the great preponderance of input to the MB main calyx is olfactory, consistent with LM data. Of the 576 microglomerular boutons in main calyx, 500 arose from olfactory PNs (87%, from 120 PNs). Of these, 20 boutons (3%) arose from 8 multiglomerular PNs. The other inputs to main calyx included 50 boutons from thermosensory PNs (9%, arising from 8 neurons); 9 boutons from other PNs (2%, arising from 5 neurons), traveling either via tracts alternative to the mALT (7 boutons from 4 PNs) or from the subesophogeal region (2 boutons from 1 putative PN; data not shown); and 17 boutons (3%) arising from a previously unknown neuron that we name MB-CP2 and describe further below. This survey located 51 out of the 52 previously described olfactory glomeruli (Grabe et al., 2015); VP4 was not located. The existence of an additional glomerulus, DL6, has been disputed (Grabe et al., 2015) and we likewise did not locate it. We also found 3 neurons arising from glomeruli VC5 or VC3l, which we could not disambiguate based on our tracing data. These glomeruli are not consistently divided in the literature, and the molecular identity of their incoming olfactory receptor neurons is not yet clear.

**Figure 4.**
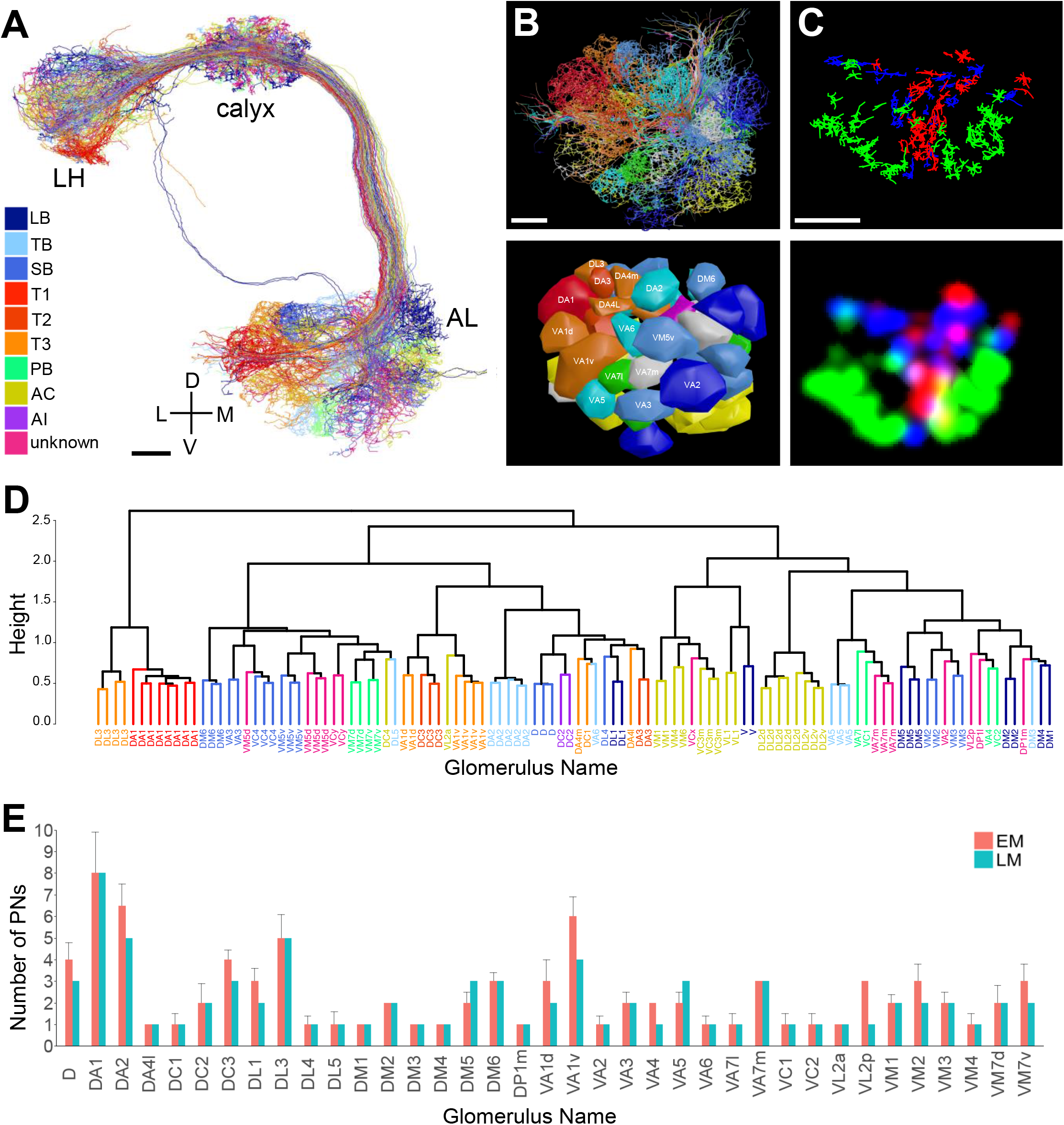
Survey of Olfactory PNs Providing Driving Input to Microglomeruli in the Main MB Calyx Agrees with LM Data. See also Table S1. (A) Frontal view of olfactory PNs on the right hemisphere. A majority of PNs extend dendrites into glomeruli in antennal lobe (AL) while their axons pass through calyx, forming *en passant* synapses with KCs, and terminate in lateral horn (LH). A few project directly to the LH via the mlALT. (B) Frontal view of reconstructed PN skeletons (upper panel) and glomerular surface models (lower panel) in AL. (C) Concentric organizations revealed in frontal-dorsal view of PN boutons in calyx. Reconstructed bouton skeletons (upper panel) and 2D projection (lower panel) of a bouton surface rendering, integrated on the Z (anterior-posterior) axis for each of 3 groups respectively. PNs from DM1, VA4, VC1, VM2 (green); DL1 and VA6 (blue); DA1, DC3, and VA1d (red). (D) Dendrogram produced by hierarchical clustering of uniglomerular olfactory PNs based on morphological similarity described by NBLAST. (E) Comparison of number of PNs per glomerulus in the EM data, versus those in Grabe et al.(2016). Colors: (A-B, D) PN colors represent sensillum type (see legend in A) for their corresponding olfactory receptor neuron (ORN) class. Color code is the same as in Couto et al. (2005) Figure 4A. Abbreviations: LB, large basiconic; TB, thin basiconic; SB, small basiconic; T1, T2, T3, trichoid sensilla; PB, maxillary palp basiconic, AC, antennal coeloconic; AI, antennal intermediate. Scale bars: ~10 μm in (A-C).

Despite these caveats, nearly all (48/52, 92%) previously described subtypes of uniglomerular olfactory PNs were unambiguously identified (Grabe et al., 2015), setting the stage for future trans-synaptic mapping of circuitry downstream of molecularly identified olfactory pathways in the fly brain. The arbors of selected subtypes formed concentric clusters in MB main calyx (Figure 4C), consistent with previous LM data (Tanaka et al., 2004). Unsupervised clustering based on NBLAST score grouped PNs of the same assigned type (Figure 4D), and in nearly all cases, the expert PN type assignments and NBLAST scores were in good agreement (Table S1). The number of PNs found to arise from each glomerulus (Figure 4E) was also in excellent agreement with recent LM data (Grabe et al., 2016).

We found that PNs arising from the same glomeruli often show much tighter clustering (Figure 5, Figure S7) than predicted from LM data pooled across multiple animals (Jefferis et al., 2007). The PN cluster at the center of the concentrically arranged arbors shown in Figure 4C (arising from DA1, DC3, and VA1d glomeruli) was also qualitatively tighter in the EM data than in LM data pooled across multiple animals (Figure 5A, bottom row). Quantification of the average distance between homotypic PNs revealed that intra-animal arbors are significantly more clustered than arbors from multi-animal LM data (Figure 5B-C). A similar result was obtained based on NBLAST score differences (Figure S7B-C). The tight clustering of EM-traced PNs suggests developmental co-fasciculation of homotypic inputs, and may bias the sampling of olfactory input by KCs (see Discussion).

**Figure 5.**
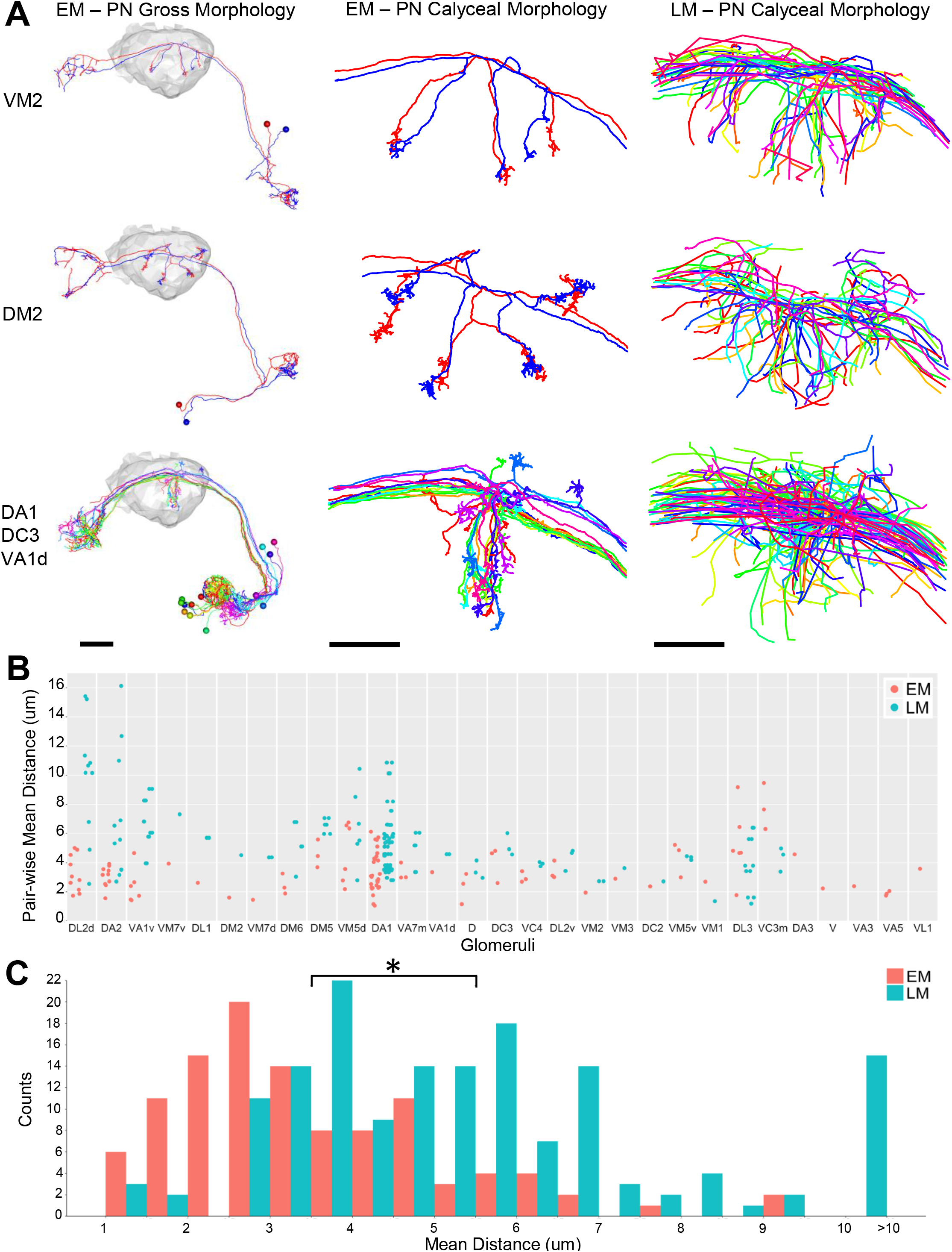
PN Arbors in Calyx Cluster More Tightly Than Previously Seen with LM Across Individuals. See also Figure S7. (A) Comparisons of PN tracings in EM and LM. Left column shows entire PNs with right calyx neuropil in gray. Middle and right columns show EM and LM PN skeletons, respectively, in calyx. (B) Pair-wise mean nearest distance for homotypic PN calyx collaterals. Glomeruli are ordered by the difference of mean distances between EM and LM PNs. Each data point represents the mean of nearest distance between the calyx collaterals of a pair of PNs from the same glomerulus. The same number of LM PNs as EM PNs is sampled from the existing database of LM neurons (Costa et al., 2016; Jefferis et al., 2007). Only glomeruli innervated by two or more PNs in the EM data are shown. (C) Histogram of all data points in (B). The total average distance for all EM PN pairs was significantly shorter than that for all LM PN pairs (3.53 ± 1.63 μm versus 5.53 ± 2.65 μm, t test p-value 1.3e-12). Scale Bars: ~20 μm in (A) left column; ~10 μm in (A), middle and right columns.

### A previously unknown cell type, MB-CP2, provides input to Kenyon cell claws

To assess the utility of the whole-brain EM dataset for characterizing previously unknown cell types, we chose to make a fuller reconstruction of one of the unidentified microglomerular inputs to the MB calyx mentioned above, which we name MB-CP2 (“Mushroom Body Calyx Pedunculus #2”; Figure 6, Movie S5), per the naming convention of Tanaka et al. (2008). We traced this neuron’s backbone to completion as well as that of its equivalent on the contralateral hemisphere (Figure 6A). The same 10 neuropil compartments were symmetrically innervated by each MB-CP2 neuron on either side of the brain (Figure 6A,F). In contrast to PNs, which receive input from olfactory receptor neurons (ORNs), MB-CP2 receives input from higher-order compartments in the protocerebrum, far from the sensory periphery (Movie S5). These include the superior medial protocerebrum (SMP), superior intermediate protocerebrum (SIP), and superior lateral protocerebrum (SLP), which are relatively little-studied compartments innervated by both sensory and motor neurons (Tschida and Bhandawat, 2015). MB-CP2 dendrites in the MB pedunculus and γ1 compartment of the MB lobes are also postsynaptic to KCs, specifically the γ (Figure 6B-C) and γd (data not shown) subtypes. In the MB main calyx, MB-CP2 provides microglomerular bouton input to all 5 olfactory KC subtypes (γ, αβc, αβs, α′β′m, and α′β′ap), but only in the anteroventral main calyx (Figure 6D-E). In the MB dorsal accessory calyx (dAC), which receives gustatory, thermosensory, and visual inputs (Kirkhart and Scott, 2015; Vogt et al., 2016; Yagi et al., 2016). MB-CP2 is presynaptic to αβp KCs throughout the entire dAC (data not shown). The two MB-CP2 neurons may therefore provide multimodal and recurrent feedback from γ KC axons to a subset of KC dendrites in the main calyx, adding to the set of known MB recurrent networks (Aso et al., 2014; Owald and Waddell, 2015).

**Figure 6.**
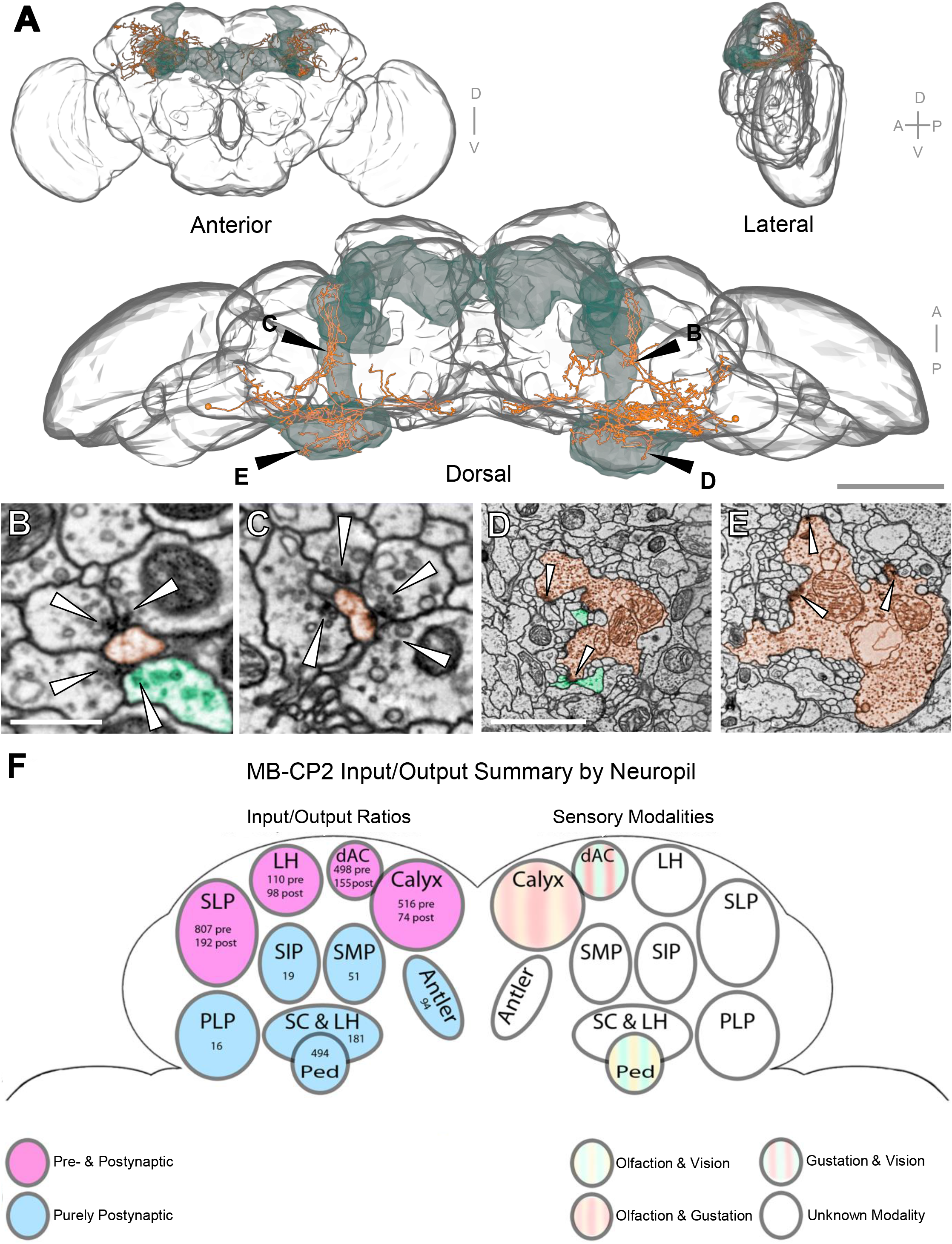
MB-CP2, a New Cell Type Providing Microglomerular Input to KC Claws. (A) 3D rendering of this neuron in both hemispheres with LM meshes of whole brain (gray) and MB (green). (B-E) TEM of synapses between MB-CP2 and KCs. MB-CP2 processes (orange); confirmed KC processes (green). (B-C) Example synapses of MB-CP2 postsynaptic to yKCs in pedunculus, right and left hemispheres, respectively. (D-E) Example synapses of MB-CP2 microglomerular organization in the main calyx, right and left hemispheres, respectively. (F) Summary schematic of MB-CP2 input and output brain regions with synaptic counts discovered thus far. This neuron innervates 10 distinct neuropils. Abbreviations: Ped, pedunculus; LH, lateral horn; dAC, dorsal accessory calyx; SC, superior clamp; PLP, posterior lateral protocerebrum; SMP, superior medial protocerebrum; SIP, superior intermediate protocerebrum; SLP, superior lateral protocerebrum. Scale Bars: 100 μm in (A), dorsal view; 500 μm in (B-C); 2 μm in (D-E).

### Identification of cell types post-synaptic to Kenyon cells in the MB calyx

Kenyon cells are presynaptic in the MB calyx, but their postsynaptic targets are unknown (Butcher et al., 2012; Christiansen et al., 2011). To identify these postsynaptic partners, we annotated all presynaptic release sites arising from 3 KCs of each subtype (γ, αβc, αβs, α′β′m, and α′β′ap) with dendrites in the main calyx (Aso et al., 2014). We then traced their postsynaptic targets to classification (Figure 7; see Methods). All KC presynaptic release sites targeted multiple postsynaptic processes. Consistent with immunohistochemical data (Christiansen et al., 2011), most (82%; Table S2) pre-synaptic release sites were in αβc-, αβs-, or γ KCs, and 87% of the release sites were distributed along KC dendrites outside of claws. Of the 15 cell types known to arborize within the MB calyx (Aso et al., 2014; Burke et al., 2012; Busch et al., 2009; Chen et al., 2012; de Haro et al., 2010; Mao and Davis, 2009; Roy et al., 2007; Tanaka et al., 2008), we found that a small subset contributes most of the postsynaptic neurites (4 subtypes contributing 75% of neurites; see Table S2). These are: the dendrites of other KCs; the APL, a wide-field inhibitory neuron that innervates the entire MB and sparsifies KC activity (Lin et al., 2014; Liu and Davis, 2009); MB-CP1, an MBON whose dendritic arbor innervates the calyx and pedunculus (Tanaka et al., 2008); and two MB-C1 neurons, a class of interneuron that innervates the calyx and lateral horn (Tanaka et al., 2008). Fourteen percent of fine postsynaptic neurites were too difficult to readily trace back to parent backbone. Intriguingly, α’β’ KCs were presynaptic only to APL and other KCs. A large fraction of KC presynaptic release sites therefore targets a specific and sparse subset of available cell types in calyx.

**Figure 7.**
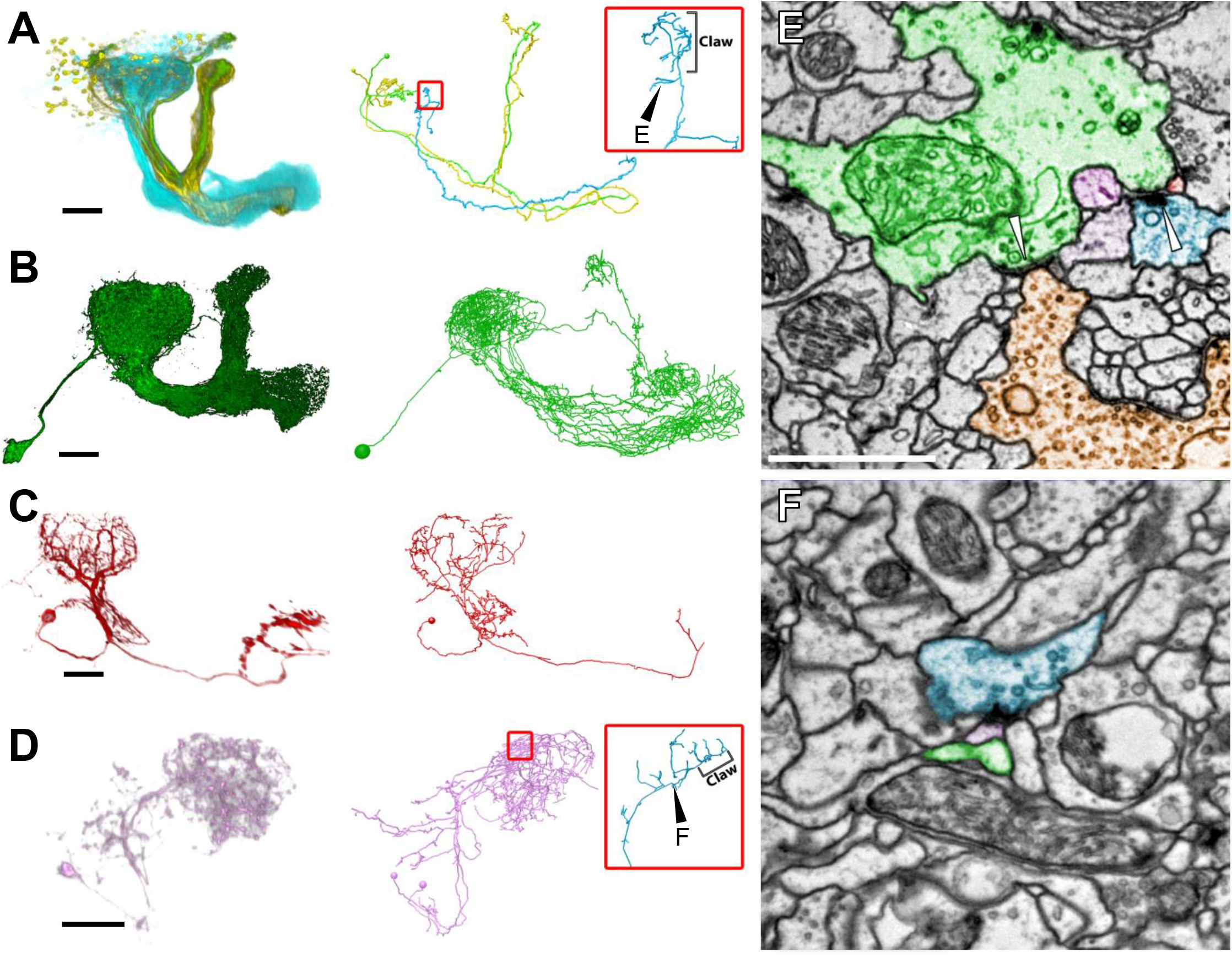
KC Presynaptic Release Sites in the MB Main Calyx Mostly Target a Small Subset of Available Partners. (A-D) Morphological comparison of LM data (left) and EM-reconstructed skeletons (right) for the same classes of neurons. (A) αβc- (green), αβs- (yellow), and γ- (cyan, blue) KCs. LM data shows the entire population for these three KC classes. EM data shows one KC of each of the three classes. Inset location indicated by the smaller red box. Inset shows the dendritic arm and claw of the γ KC that is presynaptic in (E). Black arrowhead indicates the location of the synapse in (E). Note the synapse is outside of the KC claw. (B) The APL neuron. (C) The MB- CP1 output neuron. (D) The MB-C1 putatively GABAergic interneuron. Inset location indicated by the smaller red box. Inset shows the dendritic arm and claw of the γ KC that is presynaptic in F. This KC is not shown in D for visual clarity. Black arrowhead indicates the location of the synapse in F. Note the synapse is outside of the KC claw. (E-F) TEM micrographs of KC divergent polyadic presynaptic release sites in the MB main calyx. White arrowheads indicate visible presynaptic release sites. In general the same color code is used to indicate same classes of neurons between (A-D) and (E-F). Black arrowhead (A) points to the same location in the 3D skeleton as white arrowhead points to in EM micrograph (E); same is true for black arrowhead in (D) and white arrowhead in (F). (E) The γ KC from A inset (blue) and two other γ KCs (light purple and dark purple, presynaptic release sites not visible in this section) are convergently presynaptic to the APL (green), the MB-CP1 (red), and each other at the same synaptic cleft. The APL is presynaptic to a PN (orange) in this section plane. (F) The γ KC from D inset (blue) is presynaptic via a divergent polyad to MB-C1 (pink), and the APL (green) two sections away (not visible in this section), and several additional unidentified partners. Scale Bars: ~25 μm in (A-D), 1 μm in (E-F).

## DISCUSSION

Here we contribute a complete EM volume of an adult female *Drosophila melanogaster* brain for free use by the research community. We identified PNs from nearly all the olfactory PN subtypes, and then traced PN output across two synapses – from PNs to KCs, and from KCs to their post-synaptic targets in the MB calyx – demonstrating that this dataset supports tracing of brain-spanning neural circuitry at synaptic resolution. Cell type identification of PNs was helped by software to search EM-traced neuronal arbors for matches in large-scale morphological databases. With the PN types classified, any molecularly identified olfactory pathway in the fly brain can now be mapped, which will likely aid in the determination of circuit mechanisms underlying intrinsic and learned behavioral responses to odors. Since PN odorant tuning, molecular identity, and morphology are all highly stereotyped, a great deal of information from other experiments can be directly related to the circuits mapped in this dataset. This same approach is generalizable to many other circuits underlying learned and intrinsic behaviors in this animal.

Generation of volume EM data of this scale remains technically challenging across all stages of the data pipeline. Our new generation of image acquisition hardware provided images with excellent signal-to-noise and unmatched throughput. Many further optimizations of this hardware are available. Emerging large-format, high-speed fiber-coupled cameras and direct electron detectors may achieve imaging throughput comparable to the TEMCA2, while requiring substantially lower electron dose due to their greater sensitivity (Ruskin et al., 2013). Multibeam-SEM also shows great promise (Eberle et al., 2015), as do slower but higher resolution methods such as parallel FIB-SEM imaging of slabs cut by hot knife methods (Xu et al., 2017). Low resolution EM imaging, followed by high resolution re-imaging of synaptic connectivity in selected sub-volumes, also holds promise for brain-spanning connectomics in larger animals (Hildebrand et al., 2017). Manual sectioning of long series of thin sections is not routinely replicable by most practitioners; efforts are currently underway to automate this process.

Stitching and registration of serial section mosaics at the scale of this dataset posed a significant challenge. We developed scalable software for volume reconstruction and image intensity correction, as well as a data store for managing the image transformations between raw data and any given volume reconstruction. Although the resulting registration quality is clearly sufficient for manual tracing efforts, remaining fine-scale imprecision may need to be overcome before emerging automatic segmentation methods can be fully leveraged (Arganda-Carreras et al., 2017; Beier et al., 2017; Januszewski et al., 2016). Early segmentation results with subsets of this whole-brain dataset are nonetheless promising (Funke et al., 2016).

Our analysis of the MB main calyx revealed that PNs arising from the same glomerulus often cluster more tightly in the calyx than was expected from LM data pooled across many animals. Interestingly, the intra-animal dye-filled DM6 PN pairs in Figure 6 of (Kazama and Wilson, 2009) also are tightly clustered, although this result is anecdotal. The basis of this clustering may be developmental, with sister PNs arising from the same glomerulus or neuronal lineage (Spindler and Hartenstein, 2010) tending to co-fasciculate. Tight intra-animal clustering of PNs raises the possibility that the PN to KC connectivity matrix may be biased, rather than fully random. If boutons from a given PN type are clustered tightly in the calyx, and a given KC happens to have a distribution of claws centered on that PN cluster, then the KC will have greater opportunity to receive input from that PN type. This may explain the above-chance convergence of DA1, DC3, and VA1d PNs onto postsynaptic KCs observed by Gruntman and Turner (2013). Indeed, our EM reconstructions indicate that these three PN types are tightly clustered at the center of the MB calyx, consistent with the earlier LM data of Tanaka et al. (2004). However, the most thorough examination of PN to KC connectivity to date, using partial connectivity data pooled across many animals, was consistent with a model in which the PN to KC connectivity matrix is entirely random (Caron et al., 2013). More comprehensive mapping of the KC population postsynaptic to PNs will help determine whether intra-animal biases in the PN to KC connectivity map exist, and the effect this bias may have (if any) on the overall KC sampling of olfactory input.

Our survey of input to KC claws in the MB calyx also revealed a new cell type, MB-CP2, which likely provides recurrent and multimodal input to a small fraction of KCs in the main calyx. Even in well-described brain regions, it is not uncommon for new cell types to be discovered by EM (Helmstaedter et al., 2013; Takemura et al., 2017a), or by LM in combination with increased coverage of genetic driver lines (Aso et al., 2014). The finding of a new input cell type to KC claws is also consistent with the “projection neurons innervating unknown regions of the brain” occasionally seen by Caron et al. (2013); see their Supplementary Table 1. Development of a split-GAL4 driver line for MB-CP2 would facilitate characterization of this neuron’s role in MB circuitry.

There is extensive recurrent microcircuitry between neurites within the MB calyx (Butcher et al., 2012), but the cell type identity of participating neurons has been elusive. We traced neurons postsynaptic to KC dendrites to identify their cell types, setting the stage for future interrogation of these fine-scale interactions by complementary high-resolution physiological and anatomical methods. We discovered that KC dendrites predominantly target a sparse subset of available cell types, including the wide-field inhibitory neuron APL, other KCs, the MBON MB-CP1, and MB-C1, an inhibitory neuron that innervates calyx and lateral horn. Interestingly, α’β’ KC dendrites are even more selective, targeting only the APL and other KCs. This may be related to their specific role in memory and learning; unlike other KC subtypes, α’β’ KCs are dispensable for memory retrieval (Krashes et al., 2007). Recurrent, fine-scale microcircuitry seems to be a general feature of the fly neuropil (Meinertzhagen and O’Neil, 1991; Schurmann, 2016; Takemura et al., 2017a; our unpublished observations), and identification of participating cell types will be an important initial step toward understanding microcircuit operation in many areas of the brain.

*Drosophila* exhibits a wide range of complex sensory- and memory-guided behaviors, including visual place learning, tactile-guided sequential grooming, olfactory-memory-guided courtship, escape, and vision-guided flight. The algorithms underlying behavior are implemented by neuronal circuits, and neuronal circuits are defined in large part (though certainly not entirely; Bargmann and Marder, 2013) by the synaptic connectivity between neurons. The connectome therefore is necessary to Marr’s (1982) implementation-level of analysis, and may aid in the inference of underlying algorithms as well. The dataset we share here will help establish a structural scaffold for models of circuit function in the fly.

## Author Contributions

Conceptualization, D.D.B., R.D.F., J.S.L., Z.Z.; Methodology, D.D.B., R.D.F., J.S.L., D.M., J.P., C.G.R., O.T., Z.Z.; Software, J.B., P.H., G.S.X.E.J., B.K., K.K., M.K., M.N., D.M., E.P., E.T.T., O.T., S.S.; Validation, J.S.L.; Formal Analysis, D.D.B., G.S.X.E.J., J.S.L., M.N., Z.Z.; Investigation, I.J.A., D.D.B., S.A.C-S., C.B.F., L.K., J.S.L., C.G.R., N.S., Z.Z.; Data Curation, J.B., R.D.F., J.S.L., D.M., M.N., E.P., C.G.R., S.S., Z.Z.; Writing – Original Draft, D.D.B., J.S.L., M.N., E.P., C.G.R., Z.Z; Writing – Review and Editing, D.D.B., G.S.X.E.J., B.K., N.S., J.S.L., Z.Z.; Visualization, D.D.B., S.A.C-S., J.S.L., Z.Z.,; Supervision, D.D.B., J.S.L.; Project Administration, D.D.B.; Funding Acquisition, D.D.B.

## Acknowledgements

Jeff Jordan, Bruce Bowers, and Jon Arnold (Janelia Instrumentation Design & Fabrication team); Dana Dunkelberger (Grant Scientific); Jim Mancuso (AMT Imaging): hardware design & engineering services. Brett Mensh, Scott Waddell, and Vivek Jayaraman: critical improvements to the manuscript. Wei-Ping Li (Janelia EM Facility): coating of sample grids and poststaining of serial sections. Najla Masoodpanah, Joseph Hsu, Benjamin Gorko, Emily Moore, Arynne Boyes, Jacob Ratliff, Adam John, Bailey Harrison, Adeleso Adesina, and Claire Managan: neuron tracing. Noah Nelson: graph visualization tools. Arlo Sheridan and Ben Gorko: 3D animations. Philipp Ranft and Gaia Tavosanis: literature review of cell types innervating MB calyx. Adam Heath, Marta Costa, and Philip Schlegel: glomerulus meshes, NBLAST analyses, and PN classification. David Peale: optimization of carbon coating and film casting protocol. Jon Arnold: technical figures. Tom Kazimiers and Andrew Champion: CATMAID development. Albert Cardona: CATMAID workflow advice, CATMAID development, and helpful discussions. Casey Schneider-Mizel: analysis code and helpful discussions. Tom Dolafi, Cristian Goina (Janelia Software Engineering): volume reconstruction support. Rob Lines, Ken Carlisle, Goran Ceric (Janelia Scientific Computing Systems): data and cluster management. Yoshinori Aso: KC and pre-publication MCFO PN LM data. Eyal Gruntman: helpful discussions. Howard Hughes Medical Institute: funding support for all aspects of this work. Wellcome Trust: collaborative award 203261/Z/16/Z for funding additional tracing effort. Competing interests: D.D.B., J.P. (patent holders).

## STAR METHODS

### CONTACT FOR REAGENT AND RESOURCE SHARING

Further information and requests for resources and reagents should be directed to and will be fulfilled by the Lead Contact, D.D.B.

### EXPERIMENTAL MODEL AND SUBJECT DETAILS

Multiple brains of 7 day-old [iso] *w^1118^* x [iso] Canton S G1 adult female flies were screened and one was picked for EM imaging.

### METHOD DETAILS

#### Sample preparation

Brains from 7 day-old adult [iso] *w^1118^* x [iso] Canton S G1 flies were dissected in cold fly saline (Olsen et al., 2007). The dissected brains were fixed with 2% glutaraldehyde in 0.1M sodium cacodylate for 1 hour at 4°C, followed by 1 hour at room temperature (RT). Following aldehyde fixation, the brains were rinsed 6 x 5 min with sodium cacodylate buffer at RT, 3 x 10 min incubations in 0.02M 3-amino-1,2,4-triazole (A-TRA) (De Bruijn et al., 1984) (Sigma-Aldrich) in sodium cacodylate, the last on ice, followed by post-fixation with 1% OsO_4_ in sodium cacodylate containing 0.1M A-TRA for 90 minutes on ice. The brains were then rinsed with cold sodium cacodylate buffer, allowed to warm to RT followed by deionized or Milli-Q water at RT before being stained *en bloc* with 7.5% uranyl acetate in water overnight at 4°C. Following *en bloc* staining, brains were rinsed with water at RT and then dehydrated in an ascending ethanol series to 100% ethanol, followed by 100% propylene oxide. Samples were infiltrated with EmBed 812 resin using propylene oxide to resin ratios of 2:1 and 1:2 for 30 minutes each followed by two 1-hour long incubations in 100% resin and a third 100% resin incubation overnight. Finally, samples were flat embedded between Teflon-coated glass slides and allowed to harden for 24 hours at 65-70° C.

Samples were subsequently screened for whole-brain sectioning by X-ray tomography using an Xradia XRM-510 X-ray microscope (subsequently acquired by Zeiss). Samples without obvious surface defects due to dissection, or internal defects were re-embedded in silicon rubber molds for sectioning (Fig. S3).

#### Sample supports, ultramicrotomy, and post-staining

Custom bar-coded grids made from 100 μm thick copper beryllium with a 2 × 1 mm slot, a unique serial identifier in human readable and 2-D barcode form and with fiducial markers were used to collect sections. Schematics and vendor information for the custom grids are available to non-profit research organizations upon request. Grids were prepared for picking up sections by first applying a silver/gold-color film of Pioloform (Pioloform FN, Ted Pella catalog #19244) followed by a ~8 nm layer of carbon. The Pioloform film was made thicker than normal to provide enhanced sample stability under the higher beam current necessary for rapid imaging (see below). To prepare the Pioloform film, a 600 μL aliquot of 2.05% Pioloform in dichloroethane was applied to an ethanol and hydrofluoric acid cleaned glass microscope slide (Gold Seal, Ted Pella catalog #260210) via spin coating using a Laurell WS400B-6NPP/Lite spin coater. After applying the Pioloform solution, the slide was spun for 1.4 seconds with a target speed of 8,000 rpm and an acceleration index of 255. The film was released from the slide by scribing the edges of the slide with a diamond scribe and slowly submerging the glass slide at a shallow angle into a large dish of water. The film remains floating on the surface of the water and cleaned grids were then carefully placed, bar code side down, onto the film. The film and grids were subsequently picked up from above on a 1 x 3 inch slotted and anodized aluminum slide. The anodized surface also provided a stable and reusable surface from which the grids could be cut from the surrounding support film using a heated tungsten filament. Grids were loaded onto custom 203-place stainless steel plates for carbon coating.

Carbon coating was carried out in a Denton Explorer 14 high vacuum evaporator equipped with oil diffusion pump, liquid nitrogen cold trap, and a film thickness monitor using carbon rods (Ted Pella catalog #62-132). The carbon rods were de-gassed at sub-evaporation currents (8-14 amps) prior to and immediately following sample loading. The 203-place plate was held at a 90- degree angle to the source at a distance of 10 cm during evaporation. Following a vacuum recovery period, the carbon rods were de-gassed and warmed at sub- to near-evaporation currents (8-16 amps). To avoid overheating the films, carbon was evaporated in a series of cycles (in our hands, each cycle was stopped when the deposition rate reached -0.5 Å/sec and resumed when the deposition rate returned to 0 Å/sec). Vacuum levels prior to evaporation were ~5x10^-8^ torr or better. Evaporation was carried out at 22 ± 1 amps. Carbon evaporation was halted at an indicated thickness of 70 to 80 Å and final thickness assessed after a 5 minute cool down period. Successfully prepared grid films remained perfectly flat when held within ~1 mm of a water surface (Figure S3F) whereas unsuccessful films displayed a relaxation of the film tension when held close to water (Figure S3E). Grid batches in which coatings tested did not remain flat were rejected.

Serial sections of the brain were cut with a Leica UC-6 ultramicrotome at a thickness of 35-40 nm, with periodic retrimming of the block face. Total sectioning time was ~3 weeks. Typically, 3 serial sections were collected on each of the ~2,400 custom bar-coded grids needed to collect the 7000+ sections necessary to encompass the whole brain.

Following sectioning, grids were stained in 3% aqueous uranyl acetate for 20 minutes followed by Sato’s lead (Sato, 1968) for 5 minutes, with ddH2O washes after each staining step. To facilitate the staining of ~2,400 grids, a custom Plexiglass staining device with slots to hold 100 grids at a time, loosely based on the Hiraoka (1972) device, was used.

#### Electron Microscopy

Two FEI Tecnai Spirit BioTWIN TEMs were used to image the whole fly brain series. The first, a TEMCA2 system (Figure 1C, Figure S2A), was equipped with a custom single-axis Fast Stage, vacuum extension, scintillator (5 μm Mylar on a support ring 9 ^5^/_8_ inches in diameter, coated with 10 mg fine-grained P43/cm^2^; Grant Scientific), and four Fairchild SciMOS 2051 Model F2 5.5 megapixel cameras (2560 x 2160 pixel sensor size) configured in a 2 x 2 array. The second TEM was equipped with an Autoloader (Figure 1G, S2E), a custom scintillator (6 mg fine-grain P43/cm2; Grant Scientific), and a single Fairchild SciMOS camera. In both systems, 4:1 minifying C-lenses (AMT) were mounted on the SciMOS cameras using custom lens mounts (AMT). These systems were previously described in abstract form (Robinson et al., 2016). Schematics and model files for the Fast Stage and Autoloader are available to non-profit research organizations upon request.

The Fast Stage has a single high-speed axis of motion, and is designed to interface an FEI CompuStage goniometer (Figure S2B), which provides the other degrees of freedom necessary to position a sample in the TEM. The sample holder is connected to a drive rod, which passes through a custom rolling-element bearing, vacuum sealing bellows, and a rolling-element damper (Figure 1D, S2B). The drive rod is connected to a slide-mounted encoder which provides nanometer-resolution positional feedback. It is moved linearly by a precision piezo motor (Physik Instrument cat N301K151). The custom rolling-element tip bearing provides rigid lateral support to the drive rod within the outer drive rod tube, while minimizing axial friction required to move the driven mass of the system. The custom rolling-element dampers kill vibrations of the drive rod induced by the pulsed motion of the piezo motor during moves. Without these dampers, the drive rod would vibrate for hundreds of milliseconds under the pulsed motion of a move, rendering the system unusable. With the dampers, 8-24 micron moves are reliably achieved where all vibrations are damped to less than 5 nanometers in less than 50 milliseconds (Figure S2C). The miniature vacuum bellows isolates the specimen-holding region of the device from atmospheric pressure of the operating environment. By locating the vacuum bellows just behind the O-ring in an FEI style holder, the volume needed to be evacuated after sample insertion is minimized, allowing samples to be exchanged in the same amount of time as a conventional holder.

The Autoloader GPS (Figures 1I, S2E-F) is a complete replacement for the FEI CompuStage goniometer and specimen holder, and provides all required degrees of freedom to position a specimen within the TEM column. High-speed single-axis motion is supported by the same drive mechanisms used in the Fast Stage. Other axes of motion are provided by piezo-driven and brush motors (Figure S2). The rotational angle of the sample can be changed by placing the sample grid on a rotary pre-aligner, rotating to the desired angle, and picking the sample back up again in the gripper (Movie S3). The machine vision system enabling automated handling of samples in the Autoloader recorded continuous video while operating, providing visual confirmation of proper operation and an invaluable debugging tool in the event of handling errors. To enable a continuous video stream as well as high dynamic range images suitable for image processing, the acquisition stream automatically adjusts image gain and exposure time for the required regime. These changes can be seen in Movie S3.

Image acquisition on the TEMCA2 system was performed at an indicated scope magnification of 2900x, while the single camera Autoloader equipped system operated at 4800x indicated magnification. The longer vacuum extension of the TEMCA2 system enlarged the projected image by ~1.7x, resulting in ~4 nm/pixel for both systems.

Software control of the TEMCA2 and Autoloader systems was written in LabVIEW (National Instruments). Wrapper software to interface the Fairchild SciMOS cameras with LabVIEW was written in C. Hardware triggers were used to interleave stage motion with camera frame buffer acquisition. Each camera was read out by a dedicated analysis workstation (Dell), or ‘acquisition node,’ connected via 10 Gb Ethernet to a central ‘control node’ which managed hardware triggering, stage control, region of interest (ROI) specification, mosaic preview, and user interface for hardware control. Low-latency TEM hardware control (such as beam blanking, valve operation, CompuStage control, magnification and focus adjustments, and electron beam diameter) was achieved by direct communication between LabVIEW software and the FEI dynamic-link library (DLL) files supporting the FEI Tecnai scripting environment, through the DLLs’ component object model (COM) interfaces.

Acquisition nodes measured translational drift between successive image frames in near real-time, using the NI Image analysis package (National Instruments). If drift exceeded a user-specified threshold, they were discarded and additional frames were acquired until the requested number was acquired or until a user-specified timeout was exceeded. Each acquisition node allocated three tiers of memory buffer to the image processing pipeline, to allow real-time acquisition to continue unimpeded, regardless of variations in CPU load, operating system memory management, disk performance, or network throughput. In the first tier, raw image frames were processed for drift estimation. In the second tier, sets of image frames were translated (to correct for small translations by the sample stage), summed, normalized to a background image of the scintillator, and histogram-adjusted. In the third tier, the summed and normalized images were written to disk. As images exited each of these buffers, memory was recycled so new images could be acquired and processed. Due to the rapid rate of data acquisition, multiple storage servers, each connected via 10 Gb Ethernet, were written to in round-robin fashion. Each server contained two RAID 6 volumes, and up to four servers were deployed in parallel during data acquisition. If a RAID 6 volume or a server went offline, images were written to other volumes in the available set. SSDs were installed in each acquisition node to allow an acquisition to complete in the event of total network failure during acquisition. This infrastructure was capable of supporting sustained output from the two TEMCAs and the Autoloader. No data were lost due to storage or network issues during acquisition of the whole-brain EM volume.

Autoloader control software was substantially similar to the TEMCA2 software except that it also controlled the Autoloader hardware. Autoloader-specific functionality included machine-vision-guided pick-and-place and pre-alignment of sample grids, automatic focus of the TEM, and region of interest relocation across grid picks. We also developed a user interface to let the operator define the sequence of imaging steps to be performed as well as accompanying microscope parameters for each step. All software for control of the SciMOS cameras, TEMCA2 systems, and the Autoloader is available to non-profit research organizations upon request.

For TEMCA2-imaged samples, a 16.2 nm/pixel pre-bake mosaic was acquired at 60 ms exposure time to pre-irradiate the sample and reduce specimen warping and shrinkage under high dose acquisition. The 16.2 nm/pixel mosaics were used to specify ROIs for 4 nm/pixel mosaic acquisition. The 4 nm/pixel mosaics were acquired at 35 ms exposure times. Frames were analyzed for drift in real time and 4 frames with less than 16 nm frame-to-frame drift were translated into pixel-level alignment, summed, intensity corrected, and saved. Mosaics were acquired in a boustrophedonic fashion column by column (figure S2D) running down the long axis of the 2 x 1 mm slot across the three sections such that use of the fast, piezo-driven stage axis was maximized during acquisition while slower CompuStage moves were minimized. Due to non-overlapping fields of view on TEMCA2, a two-step approach was utilized where a small stage displacement (~1900 pixels, or 7.6 μm) filled the gap between the fields of view was followed by a large displacement (~5500 pixels, or 22.0 μm) moving to a completely fresh field of view; this schema was utilized on both x and y axes with x and y steps being slightly different (5550/1950 and 5450/1850, respectively, big step/small step, in pixels). Accurate calibration of pixels per micron is essential for converting pixel distances into physical distances and allows for pixel distances to be kept constant while the conversion factor was varied depending on the indicated magnification of the microscope.

Samples are organized in the Autoloader as follows. The Autoloader holds a magazine (Figure 1L) containing 8 cassettes. Each cassette holds 64 sample grids (Figure 1F) for a total magazine capacity of 512 sample grids. The Autoloader affords random access to the individual grids, which can be retrieved, oriented, loaded into the TEM, imaged, and reliably returned to their proper address in the Autoloader. The Autoloader imaged samples in a two-pass routine where grids were returned to cassettes between acquiring pre-bake mosaics and 4 nm/pixel mosaics. The interval between imaging steps allows for the designation of ROIs for 4 nm/pixel imaging. To ensure that ROIs were accurately acquired, the Autoloader found the center of the grid slot every time a grid was loaded into the TEM column. This center point was used to align ROIs and correct for small differences in grid orientation resulting from the two-pass workflow.

The Autoloader system employed a single point autofocus routine at the center of each section to determine focus for each ROI acquired.

High-speed generation of mosaics necessitates high electron dose rate at the sample (typically ~180x the dose rate required for a 2 second exposure on Kodak 4489 film at 120 kV) to saturate the sensor wells within the short interval (35 ms in our case, vs. ~1-2s typical integration time). Pre-irradiation images of the grids were used to subdivide the samples into three ROI classes: (1) Included areas sufficiently free of substrate damage and contaminants to sustain imaging at the highest beam currents; (2) Excluded areas to be masked out of the data set entirely; (3) Borderline areas of usable but lower quality to be imaged at one tenth intensity.

Four sections (not consecutive) were lost during sectioning; and two grids, each containing 3 serial sections (3595-3597 and 6883-6885), were found to have ruptured support films after post-sectioning staining but prior to EM imaging. Sections with debris or cracks in the support film were imaged in two rounds: a high-dose, high-throughput round, excluding potentially fragile areas of a section; and a subsequent low-dose, slow exposure round, of the fragile region only. Twenty-seven sections in 9 grids ruptured toward the end of second round imaging when the low-dose electron beam hit artifacts. However, a majority (if not the entirety) of the section was already successfully imaged. In this case, although the sections were successfully imaged, the support film rupture precludes future re-imaging of these 27 sections.

#### Volume Reconstruction Pipeline

##### Overview

For each imaging acquisition system used, small step size calibration mosaics and a small reference mosaic in the same area on three reference grids were acquired. The calibration mosaics were used to calculate a correction for non-affine lens distortion for each camera in that particular acquisition system (Kaynig et al., 2010). The reference mosaics were then used to calculate the remaining affine distortion of each camera relative to all other cameras and acquisition systems, resulting in a global camera calibration model across all cameras and imaging systems involved. In the event that the imaging system configuration was modified (e.g. for camera refocusing or scintillator replacement), a new calibration mosaic was acquired, and new camera calibration models were calculated. Relational and non-relational databases were used to track image metadata and computed image transformations throughout the volume reconstruction process, and raw image data were processed using a custom-developed, highly scalable and efficient cluster-backed linear solver to stitch all section mosaics independently (Methods).

The majority of low-dose/high-dose (see Electron Microscopy) sections are acquired during a single session, without the sample being removed from the microscope. Therefore, a reliable first guess for relative positions of these layer patches is usually provided. Generally, low-dose/high-dose sections are registered in a process that takes advantage of components of the general registration pipeline above. Montages of individual acquisitions are generated and their point-matches stored. All montages sharing the same z-value (i.e. the low-dose/high-dose group of sections), together with reference neighbor “sandwich” sections are treated as a set of sections that are roughly aligned to each other as if they were all separate sections. This rough alignment is used to determine potential overlap of low-dose and high-dose areas. Tile-pairs are determined and their point-matches calculated and stored. Finally, all point-matches (within-layer, across low-dose/high-dose patches, and cross-layer to neighboring reference sections) are used to solve a linear system to determine transformation parameters for a seamless registration.

##### Migration of data

As noted above, camera images were written in a round-robin fashion across multiple high-speed RAID 6 storage servers. Mosaics selected for inclusion into the final reconstructed volume were copied to a centrally managed distributed file system at Janelia Research Campus offering high-throughput connectivity to the computational cluster as well as off-site backups. All images were checksum verified after file copy operations.

##### Stack management & relational database

We created a relational database for storing and querying metadata associated with the thousands of image mosaics and millions of acquired images. We use SQL Server 2012 for our production system and SQLite for development. Metadata required for downstream processing included: paths to image data (with checksums), stage coordinates, ROIs associated with nominal section numbers, ordering of sections and microscope configurations with associated calibrations. The input for the alignment process – a stack – can be generated with a single SQL query joining the majority of tables. The result is a list of images with their layer (z), stage coordinates (x,y), and camera configuration (for associating the correct lens correction model). The alignment process of the approximately 21 million images and associated projection of already-traced skeletons between alignment iterations is computationally expensive. To manage this we developed the Renderer toolkit (https://github.com/saalfeldlab/render), a set of image stack management tools and RESTful HTTP web services now in use in multiple additional projects. Renderer was designed in order to handle large scale (hundreds of millions) of individual records efficiently while supporting large-scale concurrent access for the stitching, section order analysis, skeleton mapping and intensity correction. Briefly, Renderer is able to quickly materialize (i.e. render) modified images for a set of transformation parameters using the mpicbg transformation library (https://github.com/axtimwalde/mpicbg). The use of the mpicbg library allows simple conversion between the Renderer database (a MongoDB instance) and TrakEM2 projects. For large scale rendering and coordinate mapping, we used Java standalone and Spark framework clients to allow it to be processed in bulk on a cluster.

##### Calibration mosaics

In our TEMCA2 system, we operate with a wider field of view than a conventional TEM which comes at the cost of individual images showing significant non-linear distortion. This distortion is the accumulation of camera lens-distortion, variation in camera mounting, and warping in the electron beam path. We compensated for this distortion using the lens-correction method available in TrakEM2 (Kaynig et al., 2010) followed by affine normalization between all distortion models. For each individual camera, we imaged a 3 x 3 mosaic of redundantly (60%) overlapping tiles of a neuropil region in one of our sample grids. This mosaic was then used to estimate a non-linear distortion correction model in TrakEM2. To compensate for the remaining affine distortion (scale and shear) of each of these camera models, we imaged a large reference montage in the neuropil region of three reference sections (to account for accidental section loss) that we then jointly aligned with TrakEM2. This way, we obtained a globally consistent camera calibration model for each individual camera. We repeated the calibration step each time an imaging system was adjusted, resulting in a set of 15 independent camera calibration models for the complete *Drosophila* brain.

##### Alignment

The image acquisition process provides partially overlapping images that are assumed to cover the entire region of interest. Image mosaics need to be stitched within each z-section plane, as well as aligned across z to produce a seamless volume. Details of the methods and documentation of actively used code are available at [https://github.com/billkarsh/Alignment_Projects/blob/master/00_DOC/method_overview.md; https://github.com/billkarsh/Alignment_Projects/blob/master/00_DOC/ptest_reference.md] and [https://github.com/khaledkhairy/EM_aligner].

Here we provide a summary. The reconstruction process consists of two steps. (1) Matching of putatively identical content between pairs of overlapping images; those matched point-pairs are stored in a table. (2) Using point-pairs to solve for linear (affine) transforms that map local image coordinates to a common stitched volume coordinate system.

##### Matching point-pairs within mosaics

Matching is first done within each of the serial sample sections (z-layers), considered independently of any other sections. Two neighboring images would match essentially perfectly except for very slight differential beam heating.

TEM stage coordinates provide useful guesses about which pairs of images have overlaps worth characterizing, as well as the expected relative transform between pair members that we can use to constrain content matching. For each prospective pair of images we first perform coarse matching using normalized FFT-based cross-correlation to obtain a best rigid transform between them: relative rotation and XY-translation. The expected constraint transform enters as a mask describing a disc of preferred XY-translations within the correlation image.

The coarse transform between image A and B is then refined using a deformable mesh as follows. Within the overlap region of A and B, the A-pixels remain at fixed coordinates. For the B-image pixels, we erect a mesh of triangles and each of the B-pixel coordinates within are translated into barycentric coordinates (functions of the triangle vertices) which are variables. The normalized cross-correlation between A and B can now be expressed as a function of mesh vertex coordinates. A gradient descent process is used to find vertex positions that optimize correlation.

The reported point-pairs linking A to B are derived from the triangles of the mesh. Image-point A is defined as the centroid of a given mesh triangle prior to optimization. Its corresponding B-image point is obtained by calculating the affine transform that takes the triangle to its optimized counterpart, and applying that to the A-centroid.

##### Matching point-pairs across layers

Since the layers are nominally 40 nm thick and neural processes propagate through tissue at all possible angles, content in adjacent layers is grossly similar but isn’t a precise match. Nevertheless, content-based matching as described above for same-layer image pairs (FFTs followed by deformable mesh optimization) works very well if combined again with expected pair-pair transforms for which we have high confidence.

First we match whole layers to each other: For each layer, individually, we collect the reported in-layer point matches and solve for its set of affine transform parameters that register that layer’s 2D images to form a so-called montage. These data are used to render the layer at a reduced scale (~20X) to an image that we call the “montage scape”. Scale reduction allows the problem to fit comfortably in RAM, reduces computation time, and most importantly, emphasizes larger size tissue features such as large neurites running parallel to the z-axis, which vary much more slowly as a function of z than neuropil. Each pair of montage scapes is matched by FFT cross-correlation at a series of angles and the best correspondence is determined. This is followed by manual inspection using TrakEM2 (Cardona et al., 2012) to verify this rough alignment.

##### Aligning Section Montages and Section-order Correction

For larger volumes, we implemented a fully automated procedure for whole-layer matching. SIFT features are extracted from section montages, and point-correspondences are determined for all pairs of sections within a range of expected ordering mistakes (in our case within 100 sections). We then use the number of point-correspondences between two sections as a surrogate for their inverse relative distance and identify the shortest possible path to visit all sections, resulting in an ordered series (Hanslovsky et al., 2017). Then, a regularized linear system is solved to calculate an affine transformation for each section that roughly aligns the volume.

With a given pair of layers now coarsely aligned, we subdivide each layer into an array of ‘blocks’ (~10 x 10 neighborhoods of image tiles). We again step angles and calculate FFT cross-correlation, this time on pairs of corresponding blocks to find the best block-block transforms. As a result we know which images within the blocks pair with each other and what their relative transform ought to be. Again, we subdivide each image into local regions, estimate point correspondences using FFT-based cross-correlation, and collected these correspondences in a database.

##### Solving the volume

With the full set of point pairs tabulated, each image is typically connected to several of its neighbors. We then construct a system of equations requiring that, under the sought affine parameter set that defines each image transformation, point-pairs should map to the same global point in the reconstructed volume. To avoid spurious deformation and volume shrinkage, the equation system is regularized to a roughly aligned volume. This roughly aligned volume depends on individual montages that were in turn regularized to a rigid model approximation that is independently estimated. The full system constitutes a large linear sparse matrix problem, whose solution provides the globally optimal transformation for all images simultaneously.

##### Sources of error

- Wrong (low-quality) point-pairs: These may occur due to the self-similarity of nominally good quality neural EM images. Errors are even more likely in tissue regions that are substantially devoid of neurons or texture, such as the lumen of the esophagus, or along the outer boundary of the sample where tissue is sparse or even absent from several image tiles. To address this error we employ (a) auxiliary contextual information about the likely transform between any two images that constrains matching derived from local image content alone, and (b) we impose a strict point-matching filter using Random Sample Consensus (RANSAC); (Fischler and Bolles, 1981) to separate true correspondences that behave consistently with respect to an affine transform up to a maximal correspondence displacement (Saalfeld et al., 2012).
- Missing point-matches: In some cases tissue damage, contamination or folds within a section lead to a lack of point-matches in a smaller region within the volume. This is most prominent when searching for point-matches across z. We address this issue by extending the point-match search beyond immediate neighbor sections.

##### Render (Image Intensity Correction)

During iterative volume reconstruction, gradient-domain processing is used to remove seams in two dimensions. A target gradient field is constructed by computing the gradient field of the input mosaic and zeroing out seam-crossing gradients. Then, a least-squares system is solved to find the new image whose gradients best fit the target field. In addition, low-frequency modulation is removed by computing the windowed average of adjacent mosaics and replacing the low-frequency components of an input mosaic with the low-frequency components of the average. We anticipate that future work will allow 3D processing of the whole-brain image volume (Kazhdan et al., 2015), reducing or eliminating section-to-section variations in intensity.

##### Projection of arbor tracing across alignments

With each new alignment, the CATMAID PostgreSQL database containing all neuronal skeleton coordinates (Schneider-Mizell et al., 2016) is dumped to retrieve their “world” coordinates (coordinates representing their physical location in the brain). Each of these world coordinates is then inversely transformed using the Renderer service (see “Stack management & relational database” section) to a set of “local” coordinates detailing the source tile visible at that location and the relative location within. The local coordinates are projected back into world coordinates using the new alignment’s transformations. The updated coordinates are then applied to a new copy of the database.

#### Neuron Tracing

Neuron reconstructions are based on manual annotation of neuronal arbors from image stacks in CATMAID (http://www.catmaid.org) as described in (Schneider-Mizell et al., 2016). All neurons included in analyses are reconstructed by at least 2 team members, an initial tracer and a subsequent proofreader who corroborates the tracer’s work. In the event that any tracer or proofreader encounters ambiguous features (neural processes or synapses that are not identifiable with 100% confidence), they consult other tracers and proofreaders to determine the validity of said features, climbing the experience ladder up to expert tracers as needed. If any feature remains ambiguous after scrutiny by an expert tracer, then said feature is not included in the neural reconstruction and/or flagged to be excluded from analyses. During the proofreading phase, the proofreader and tracer iteratively consult each other until each neuron is deemed complete per the specific tracing protocol to which it belongs. An assignment of completion does not necessarily entail that an entire neuron’s processes and synapses have been reconstructed (see “Tracing to Classification” and “Tracing to Completion” sections below).

The criteria to identify a chemical synapse include at least 3 of the 4 following features, with the first as an absolute requirement: 1) an active zone with vesicles; 2) presynaptic specializations such as a ribbon or T-bar with or without a platform; 3) synaptic clefts; 4) postsynaptic membrane specializations such as postsynaptic densities (PSDs). In flies, PSDs are variable, clearer at postsynaptic sites of KCs in a micro-glomerulus but often subtle, unclear, or absent in other atypical synaptic contacts (Prokop and Meinertzhagen, 2006). In the absence of clear PSDs, all cells that are immediately apposed across a clearly visible synaptic cleft are marked as postsynaptic. We did not attempt to identify electrical synapses (gap junctions), since they are unlikely to be resolved at the 4 nm x-y pixel size of this data set.

##### Tracing to Classification

Often only reconstruction of backbone (e.g. microtubule-containing ‘backbone’ neurites, (Schneider-Mizell et al., 2016) or gross morphology is needed to classify a neuron based on expert identification or NBLAST-based neuron searching against an existing LM dataset. If either approach fails to find a match (as in the case of MB-CP2 in our study), the neuron may be deemed a new cell type. Neurons traced to classification are at a minimum skeletonized, with or without synapses, to the point at which their gross morphologies (or backbone skeletons) unambiguously recapitulate that observed by LM for a given cell class, or are unambiguously deemed as a new cell type not previously observed in all LM database from NBLAST neuron morphology search and/or multiple experts.

##### Tracing to Completion

All steps for tracing to classification were completed. Additionally, every identifiable process and every identifiable synapse is traced within the data set.

##### Multiverse

Three teams each comprising 2 members, 1 tracer and 1 proofreader, reconstructed the same KC fragment to completion in tracing environments blinded to each other. In the tracing phase, the tracer had access to the proofreader for consult and verification. During the proofreading phase the proofreader had access to the tracer for consult and verification. When complete the reconstructions were merged into a single viewing environment for comparison (Figure S6).

##### Tracing of Projection Neurons

Three protocols were used to reconstruct olfactory projection neurons (PNs) on the right side of the brain: **1)** putative PN boutons presynaptic to all traced claws of ~300 KCs as part of a separate ongoing effect of KC reconstructions (data not shown) were seeded and traced to classification. **2)** A seed section at the posterolateral bend of the mALT, proximal to MB calyx, was selected and all neurons not found via protocol 1 were traced first directly toward the calyx. Neurons that innervated calyx were traced to classification, whereas those that bypassed calyx were halted. **3)** A thorough visual survey of the calyx was conducted to ensure that all microglomerular structures had been identified and the untraced boutons within these microglomeruli were seeded with single skeleton nodes then traced to classification.

Classification of olfactory glomeruli in AL followed that of Grabe et al. (2015), except that VC3l and VC3m glomeruli were treated as separate glomeruli (Chou et al., 2010; Silbering et al., 2011). Following Grabe et al. (2015) and Yu et al. (2010), VM6 and VP1 were combined into a single glomerulus due to morphological ambiguities, which we label as VM6 in this work.

##### Delimitation of Boutons in Projection Neurons

Projection neuron axonal boutons in the calyx were identified by varicosities containing arrays of presynaptic active zones each apposed to many postsynaptic processes (Figure S1A). Skeleton nodes at the varicosity/intervaricosity borders were tagged as “bouton borders” such that they contained all synapses inside each varicosity.

##### Kenyon Cells and their Calyceal Postsynaptic Partners

Three KCs from each of the KC classes that innervate main calyx (γ, αβc, αβs, α’β’m, and α’β’ap) were selected from a larger set of several hundred KCs already traced to classification as part of an ongoing study. All neurons postsynaptic to every presynaptic release site of the 15 KCs in the PN bouton-containing portion of the calyx (namely, postsynaptic partners in the calyx) were enumerated and traced to classification. Postsynaptic partners to low order KC dendrites were not traced unless these dendrites occupied the PN bouton-containing portion of the main calyx.

##### MB-CP2

The 2 MB-CP2 neurons were traced to classification per the “Tracing to Classification” section above. Additionally, samples of their synapses were traced within each neuropil they innervate. More synapses were traced for the right hemisphere neuron than the left hemisphere, as the left hemisphere neuron was traced to recapitulate the morphology and synapses observed in the right hemisphere.

#### Neuronal Informatics

##### Electron-Light Microscopy tools ELM

ELM provides a user interface to manually define a three-dimensional warp field between a light microscopy data set and the whole-brain EM dataset by specifying corresponding landmark points. It was built on top of the BigWarp Fiji plugin (Bogovic et al., 2016), which in turn was built on top of the BigDataViewer plugin (Pietzsch et al., 2015) for FIJI (Schindelin et al., 2012). ELM is aware of standard compartment boundary models available for the template fly brains and provides hotkeys to view the labels for these compartments; to go between coordinates in ELM and the EM dataset as viewed in CATMAID; and to go from a CATMAID URL to the corresponding point in ELM. ELM is available at https://github.com/saalfeldlab/elm.

##### Transforming data between EM and light microscopy templates: elmr

elmr (https://github.com/jefferis/elmr) is a package written in R (http://www.r-project.org) to facilitate bidirectional transfer of 3D data between adult brain EM and light level data.

##### Neuropil surface models

Previously defined surface models of the whole fly brain and MB calyx (Ito et al., 2014; Manton et al., 2014), based on the same template brain as the virtualfybrain.org project (https://github.com/VirtualFlyBrain/DrosAdultBRAINdomains), were transformed to the EM volume using elmr. The AL glomerulus meshes were generated in Blender (www.blender.org) from EM-reconstructed skeletons of PN dendrites and olfactory receptor neuron termini (Schlegel et al., 2016).

### QUANTIFICATION AND STATISTICAL ANALYSIS

#### Comparison of signal-to-noise between volume EM datasets

Determining the signal to noise ratio (S/N) of biological images is in general a subjective task, due to its variance under non-linear transformations (Erdogmus et al., 2004). As users of this data will likely care about biological structures, the determination of S/N should account for this, considering only the level of signal of these structures and not of things such as staining or cutting artifacts. The problem of S/N determination has been thoroughly treated in the case of super-resolution imaging where these ambiguities don’t exist (for a review, see Lambert and Waters, 2016; see also Supplementary Note 1 in Li et al., 2015), but as yet there are no universally accepted, automated techniques to calculate the S/N in individual images where signal is dense in both spatial and frequency spaces, such as EM data of brain neuropil.

We present two measures of S/N here, an automated measure which avoids user biases, but can include some signal in noise and background calculations (feature based signal-to-noise ratios) and a simple technique which gives more precise S/Ns but is prone to bias (the cell-edge technique) which we use to verify the feature based signal-to-noise calculation. We apply these techniques to a range of publicly available data in order to evaluate the TEMCA2 method, sample images from each dataset are shown in Figure S4A.

In both cases we assume that noise is additive (and is independent of the magnitude of the signal) and symmetric. Such an assumption is likely false (e.g. electron shot noise is Poissonian and not symmetric at low numbers), however, such impacts are likely small based on manual examination of images and we assume the impacts of such an assumption are the same for all techniques. Such assumptions may fail however at very low signals where CCD and shot noise dominates, or at high signals, where processes such as non-linearity in CCD absorption become important.

##### Feature based signal-to-noise ratios

Fundamentally a signal-to-noise calculation of an image involves a calculation of the background level, the variation in this background level (which is assumed to be due to noise) and the calculation of the difference between the regions of interest and this background. Detecting what these regions are provides a challenge in EM data where images may not have clear background regions and where noise is contributed to through sample preparation.

In order to measure the S/N we assume that in any given image, the structures of interest provide the majority of features above the noise. That is, most structures present are biological in nature, rather than artifacts of sample preparation. Therefore with this assumption, it can be further assumed that key-points detected by feature detection algorithms will disproportionately fall on the regions of interest.

Given that animal cells and structures therein tend to be “blobby” due to hydrostatic processes (Jiang and Sun, 2013), we use a blob-detection algorithm (which compares areas of interest, c.f. edge or corner detection) to identify areas of interest. We use the SURF algorithm (Bay et al., 2008), though SIFT (Lowe, 2004), BRIEF (Calonder et al., 2010) or other detection algorithms should provide enough feature locations to produce similar results (see for example (Kashif et al., 2016) and references therein).

Following the above, the variation in intensity of an image, *I*, in the local region of many feature points is likely to be mostly due to signal, and the variation in intensity nearby few (or no) feature points will be dominated by noise. The determination of such regions is done by generating an array of equal size to the original image and for each element, setting it to one if there is a feature in the corresponding element of the image. This array is then convolved with a Gaussian of width *n*, where *n* is chosen to maximize the SNR in a random selection of five images from each sample in an effort to avoid bias between samples.

To select a region dominated by noise we then shuffle this array before sorting it (to avoid biases in sorting algorithms) and take the lowest point. We then block out a region 2*n* square and resort the array nine more times (for a total of ten selected regions), forming the set of points *p*_low_. We likewise perform a selection for the points of maximum variation (*p*_high_). See Figure S4B for an illustration of the entire process.

To determine the level of noise, we first generate a copy of the image to which a three pixel median filter has been applied. We then subtract this median image from the original to generate a noise dominated image, *I*’. At each of the minimum feature points (where noise is most dominant); the standard deviation of this image is taken over a three pixel square neighborhood. The level of noise, *N*, is then calculated as the mean of these standard deviations, i.e.,

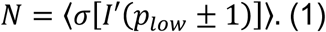

The background level of the image, *B*, is determined by taking the mean of these noise-dominated regions (again taking a mean over the three pixel neighborhood), following on from the assumption of symmetric noise, giving

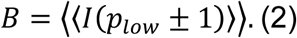

The level of signal is then taken to be the mean of the (absolute) difference of the mean of these three pixel neighborhoods around *p*_high_, and the background. This results in the SNR being given by

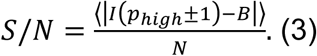

As most images lack large areas that consist of only resin, this simple background selection is not perfect, as such the S/Ns generated should be considered lower limits in most cases. We show the S/N as a function of the acquisition rate for a variety of EM techniques in figure 2C.

This measure works reasonably well when combining voxels producing S/Ns within 20% (1.5 dB) of the expected based on additive Gaussian noise (Figure S4C), although the ATUM data of Kasthuri et al. (2011) increases by more than others, a possible sign of their voxels (3x3 nm in X-Y) under-sampling biological features. This method produces the expected increase when scaling down images producing equivalent normalized S/Ns (Figure S4C). Increasing the size of images also increases the S/N, but this is due to the generation of new pixels with similar values to old ones inside the regions considered for noise due to the fact that creation of these new pixels functions as a pseudo-low pass filter. As expected this measure reports larger S/N values when Gaussian blurring is applied (as noise is disproportionately removed when a low pass filter is applied) (Figure S4C). In images generated by super resolution techniques therefore, this method may be inappropriate and should be modified to, for example, use distance based regions rather than pixel based regions.

##### Cell-membrane signal-to-noise

Although the feature based signal-to-noise measure avoids many human biases in the selection of regions used to calculate background and signal levels, it unfortunately can often incorporate biological structure (our signal of interest) into these calculations.

We therefore introduce a complementary measure to compare the S/N of biological EM data and verify that the feature based signal-to-noise calculation is valid. At its heart, this is simply a comparison between the signal level at a cell edge and the background nearby, taken at multiple points within an image.

This is achieved by a user creating a line inside a random 100×100 pixel region which contains only resin and, ideally nearby, a line which covers only a stained cell boundary. Pixels along these lines (as selected by Bresenham’s line algorithm (Bresenham, 1965)) are considered to be background or signal respectively. After selection of a background and signal line within each region, another 100×100 pixel region is chosen, until twenty lines in total (ten background, ten signal) are selected, skipping a region if there is not a suitable region in which to select both.

The noise is considered to be the standard deviation of the pixel intensities across all background intensities, and the background level the mean. The signal value is considered to be a mean of the signal pixels.

We show an example of the selection process in Figure S4D-E. Signal-to-noise ratios found via this method Figure S4E, are found to be within 10% of that found via the feature method, suggesting the former may be used for a fast, bias-free, comparison between methods.

#### Analysis of neuronal geometry

Data analysis was conducted using custom packages developed in R. We imported EM skeleton data from the CATMAID tracing environment using rcatmaid (https://github.com/jefferis/rcatmaid). Section thickness in CATMAID was specified to be 35 nm; all analyses of skeleton geometry therefore use this value. For qualitative and quantitative comparison with LM neurons, the EM skeletons were transformed into coordinate spaces of various LM template brains using elmr based on landmark pairs defined with ELM (see above). The R NeuroAnatomy Toolbox package (nat, https://github.com/jefferis/nat) was used for geometric computations, 3D visualization of neuronal skeletons and surface models.

##### NBLAST neuron search for Projection Neurons

The EM skeletons of PNs were transformed into the FCWB template brain space for NBLAST neuron search against the ~400 LM PNs previously classified in the FlyCircuit dataset by glomerulus (Costa et al., 2016). This is enabled by a single elmr function nblast_fafb. The search functionality is built on the nat.nblast package (https://github.com/jefferislab/nat.nblast) and uses data distributed with the flycircuit package (https://github.com/jefferis/flycircuit), both of which are installed with elmr. Only PNs whose candidate glomerular types exist in the FlyCircuit dataset are used. Since EM-reconstructed PNs often have many additional fine processes compared with their LM counterparts, EM skeletons were used as NBLAST targets rather than queries, in reverse to conventional NBLAST option. For each PN in EM, the top 5 hits of LM neurons and their NBLAST scores are tabulated to aid and/or confirm expert glomerular identification of PNs. Further details of the NBLAST neuron search, the associated LM data, and an online web-app for on-the-fly NBLAST queries are available at http://jefferislab.org/si/nblast.

##### NBLAST clustering for PNs

Pair-wise all by all NBLAST scores were computed for all uniglomerular PNs (nat.nblast function nblast_allbyall) after transformation into the JFRC2 template brain (Jenett et al., 2012) space using elmr. We used unsupervised hierarchical clustering with Ward’s method based on the NBLAST scores (nat function nhclust). The unsquared Euclidean distance, rather than the default square of the Euclidean distance, is used as the Y axis for dendrograms.

##### Analysis and renderings of PN Arbors in Calyx

We wished to quantify homotypic physical clustering of PNs in EM versus LM data. In summary, we randomly selected the same number of LM PNs as EM PNs from an existing LM database, subsetted the calyx arbors of the PNs with a calyx bounding box, and computed pair-wise geometric measures (mean nearest distance and NBLAST scores). Mean nearest distance quantifies physical co-location of arbors while NBLAST scores measure morphological similarity for a given pair of neurons. Details are as follows.

LM datasets (Chiang et al., 2011) of PNs previously registered to a common template brain (Costa et al., 2016) were used for comparisons with EM PNs. From the LM dataset we first determined which glomeruli had multiple EM and LM tracings available. One glomerulus, DA3, was excluded because in LM data DA3 has *en passant* collaterals that do not enter the MB calyx (Jefferis et al., 2007). We then selected a random set of LM skeletons so that we had the same number of LM and EM skeletons for each glomerulus.

Both EM and LM PNs were transformed onto a common template brain, the JFRC2 template used by virtualflybrain.org (Manton et al., 2014), and resampled with a 1 μm interval to ensure uniform representation of skeletons. PN collaterals in the calyx were obtained by two steps: 1) subset the skeletons with a bounding box defined by the right-side calyx surface model from the neuropil segmentation generated and used by the Virtual Fly Brain (Ito et al., 2014; Manton et al., 2014; Milyaev et al., 2012); 2) remove the backbone of each PNs so only the *en passant* collaterals entering the calyx are used for distance calculation.

For each glomerulus, geometric measures (mean nearest distance and NBLAST scores) were computed for each pair-wise combination of PNs of the same type. For mean nearest distance, we iterated over each node of the query neuron to find the nearest node in the target neuron, measured the Euclidean distance, and calculate the mean distance for all nodes in the query neurons (forward distance). When the query neuron is significantly larger than target neuron, artifactually long nearest distance can be introduced by end points in the larger neuron being far away from closest nodes from the smaller neuron. To address this issue, we calculated the same mean nearest distance with the query neuron and target neuron in reverse (reverse distance) and picked only the smaller of the forward and reverse distances. To quantify morphological similarity, NBLAST scores were computed for PN arbors in calyx in a similar pair-wise manner. The distributions of all intra-glomerular pairwise mean distances and NBLAST scores of PNs were plotted, and for both measures the difference between EM and LM population data were analyzed with a Student t-test.

To visualize only calyx arbors of PNs, a calyx bounding box as defined by calyx neuropil segmentation in the Virtual Fly Brain template brain (JFRC2) was used to subset both EM and LM PN skeletons. Boutons of PNs in calyx are delimited by tagged nodes bordering bouton blocks of the skeletons (See Neuron Tracing). Only skeletons in PN boutons are kept for rendering concentric organizations of boutons in calyx. For all tracing visualizations, linear interpolation of neighbouring skeleton nodes was applied to smoothen artifactual spikes in neuron tracing due to registration or alignment errors.

### DATA AND SOFTWARE AVAILABILITY

All files and movies are available through the following website: http://www.temca2data.org

### ADDITIONAL RESOURCES

Access to the full adult fly brain data set is available at: http://www.temca2data.org

Analysis code is available at: https://github.com/dbock/bocklab_public/tree/master/temca2data

## Supplemental Figure and Table Legends

**Figure S1.**
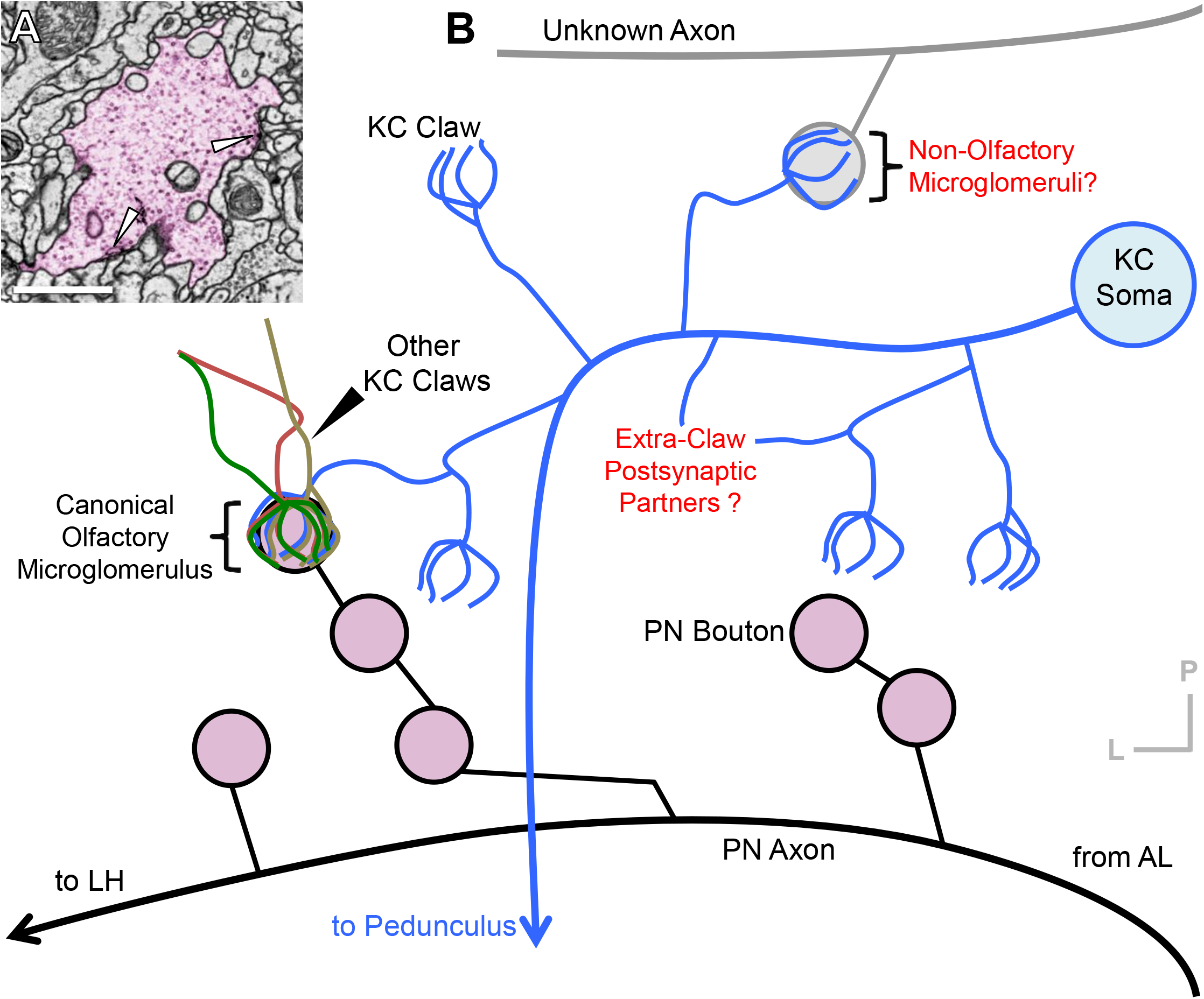
Neuronal Architecture of the MB Calyx. Related to Figure 1. (A) TEM micrograph of a calycal microglomerulus. A canonical olfactory PN axonal varicosity (bouton) is presynaptic to several KC dendritic claws. This architecture is collectively referred to as a microglomerulus. The PN terminal (pink) provides input to KC neurites at synaptic sites. Arrows: presynaptic release sites. (B) Schematic of microglomerular inputs to KCs in calyx of *Drosophila*. The PN axons from AL extend collaterals into the calyx and form boutons containing synapses to claw-shaped dendrites from several KCs. The complete composition of cell types that provide driving inputs in microglomerular form in the calyx is unknown, as is the distribution of other KC inputs outside of claws. KCs have been shown to form presynaptic release sites in the calyx most of which, but not all, are outside of claws, and the complete postsynaptic partner cohort is unknown. Scale Bar: 1 μm in (A).

**Figure S2.**
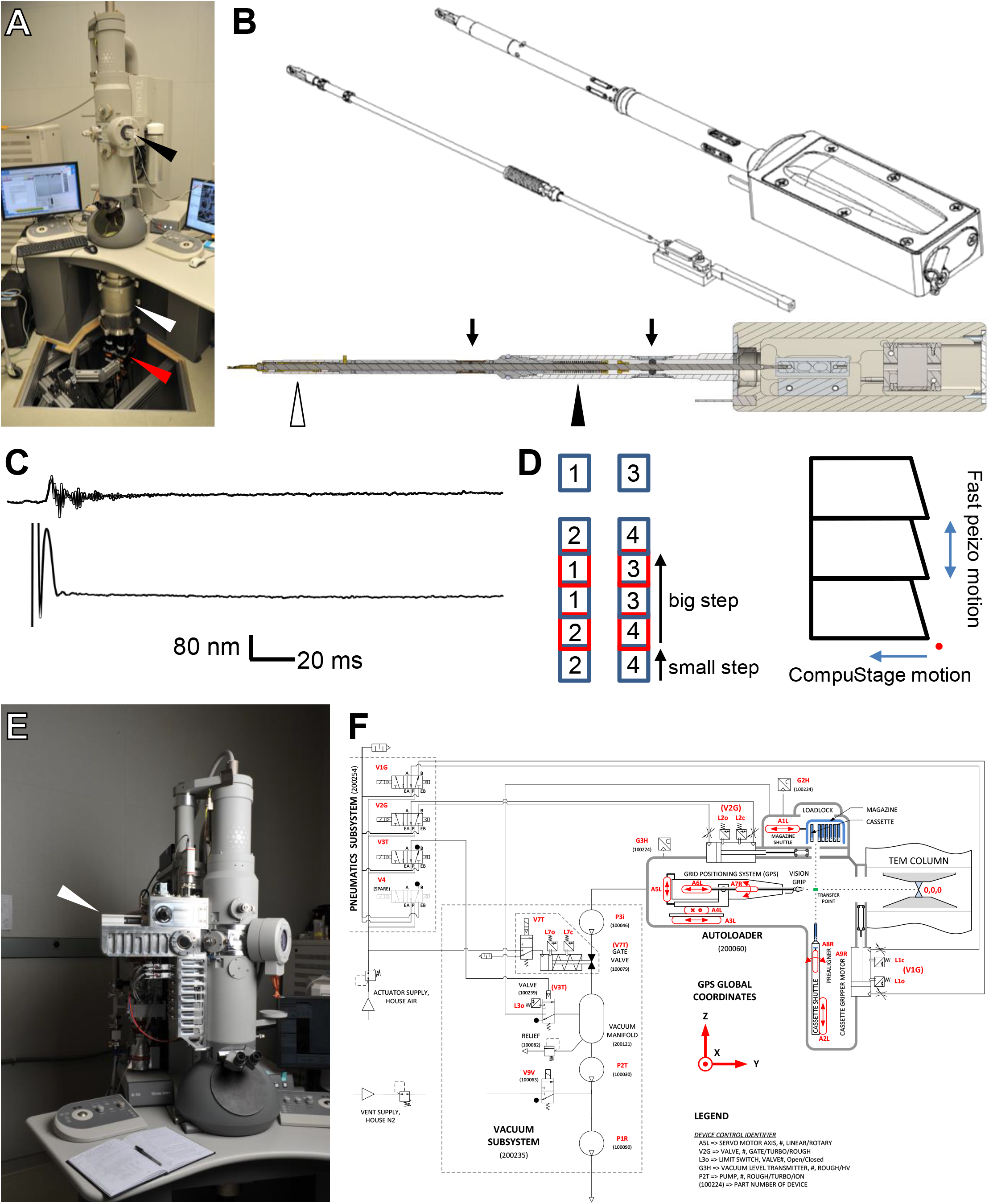
Fast Stage Step and Settle, Overview of Details. Related to Figure 1. (A) TEMCA equipped with Fast Stage. Arrowheads: Fast Stage (black); elongated vacuum chamber (white); 2 x 2 camera array (red). (B) Upper panel: Schematic of Fast Stage showing the locations of bearings, dampers and vacuum bellows. Left: driven mass; Right: exterior view. Lower panel: Cut away of Fast Stage. Arrows: rolling element damper locations (black arrows); vacuum bellows (black arrowhead); rolling-element ‘tip’ bearing (white arrowhead). (C) Plot of Fast Stage motion over time following an 8 μm move. (D) Schematic of Fast Stage stepping pattern. Left: small step/ big step schematic. Numbers are camera numbers. Right: Scanning axes and stages. Red point is origin of scanning. (E) Autoloader (white arrowhead) mounted to an accessory port on FEI Tecnai Spirit BioTWIN. (F) Schematic of the Autoloader system diagraming motor positions and movement axes as well as vacuum and pneumatic subsystems.

**Movie S1. Fast Stage vs. FEI CompuStage. Related to Figure 1.**

Movie showing the acquisition of 2 fields of view using the FEI CompuStage (left) as compared to the custom FastStage acquiring 17 fields of view (right) in the same time.

Scale Bar: 1 μm.

**Movie S2. Autoloader Cutaway. Related to Figure 1.**

CAD movie of Autoloader detailing actions to retrieve and image a sample.

**Movie S3. Autoloader Pick-and-Place. Related to Figure 1**.

Movie of Autoloader pick-and-place routine. The Autoloader locates the grid within the cassette, moves to a pre-pick location, confirms positioning, picks grid from cassette in a two step process with positioning assessments during the process, moves to the aligner, assesses the rotational angle of the grid, if necessary places the grid on the aligner and aligns the grid, retrieves the grid from the aligner, and inserts into the TEM column (insertion to column not shown). Following imaging, the Autoloader locates the correct cassette pocket, confirms positioning, replaces the grid in the cassette, and confirms that the grid is correctly located within the cassette. Changes in image quality indicate a change in camera frame rate. High quality images are used for positional assessments; lower quality images are used for diagnostic purposes.

**Figure S3.**
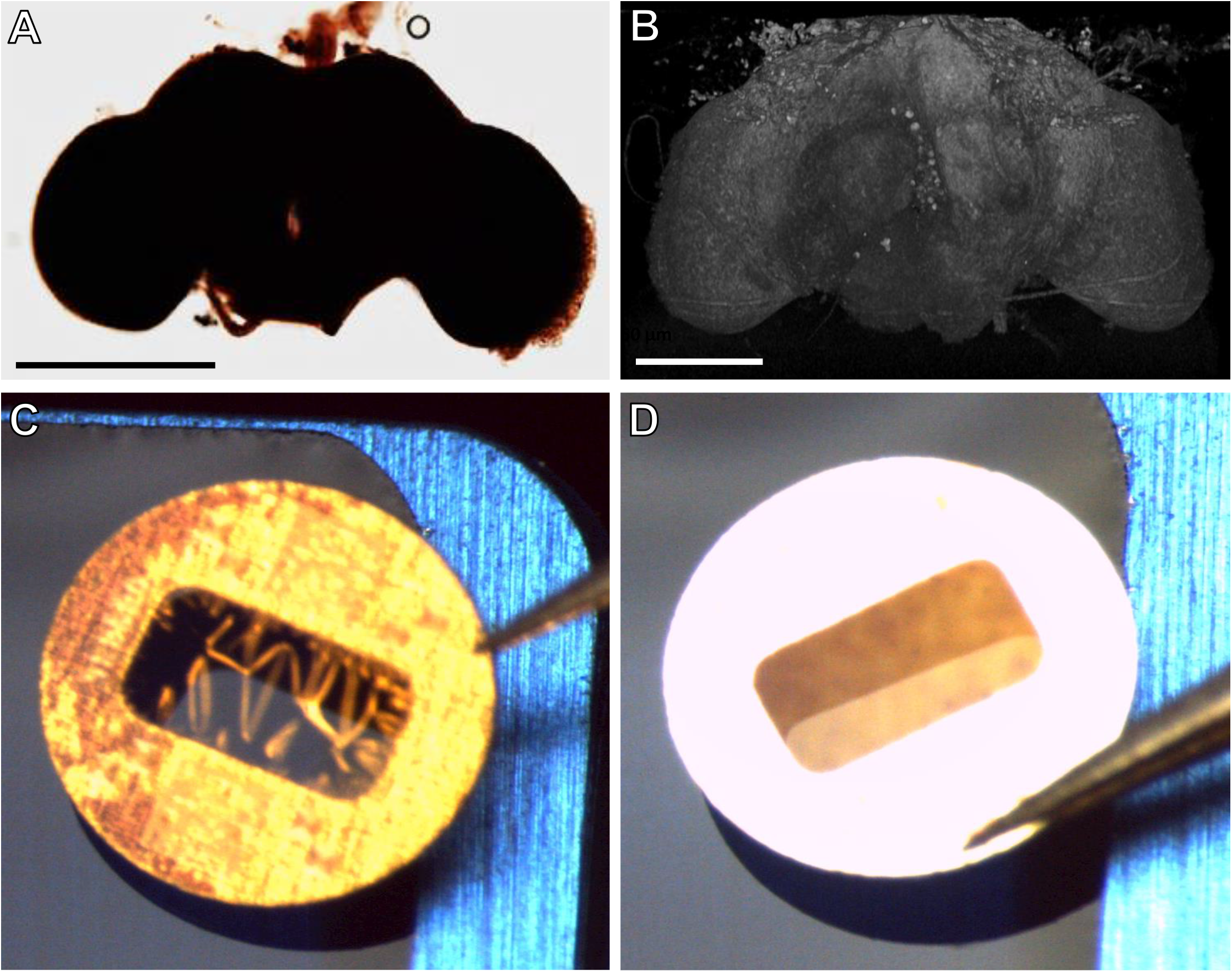
Sample Preparation for Electron Microscopy. Related to Figure 1. (A) *D. melanogaster* brain following sample preparation. (B) 3D volumetric rendering of X-ray tomogram data from embedded *D. melanogaster* brain. (C) Sample support test showing a failed result with wrinkling of the support film on 3mm grid. (D) Sample support test showing a successful result with no wrinkling or relaxation of the support film evident. Scale Bars: 250 μm in (A-B).

**Figure S4.**
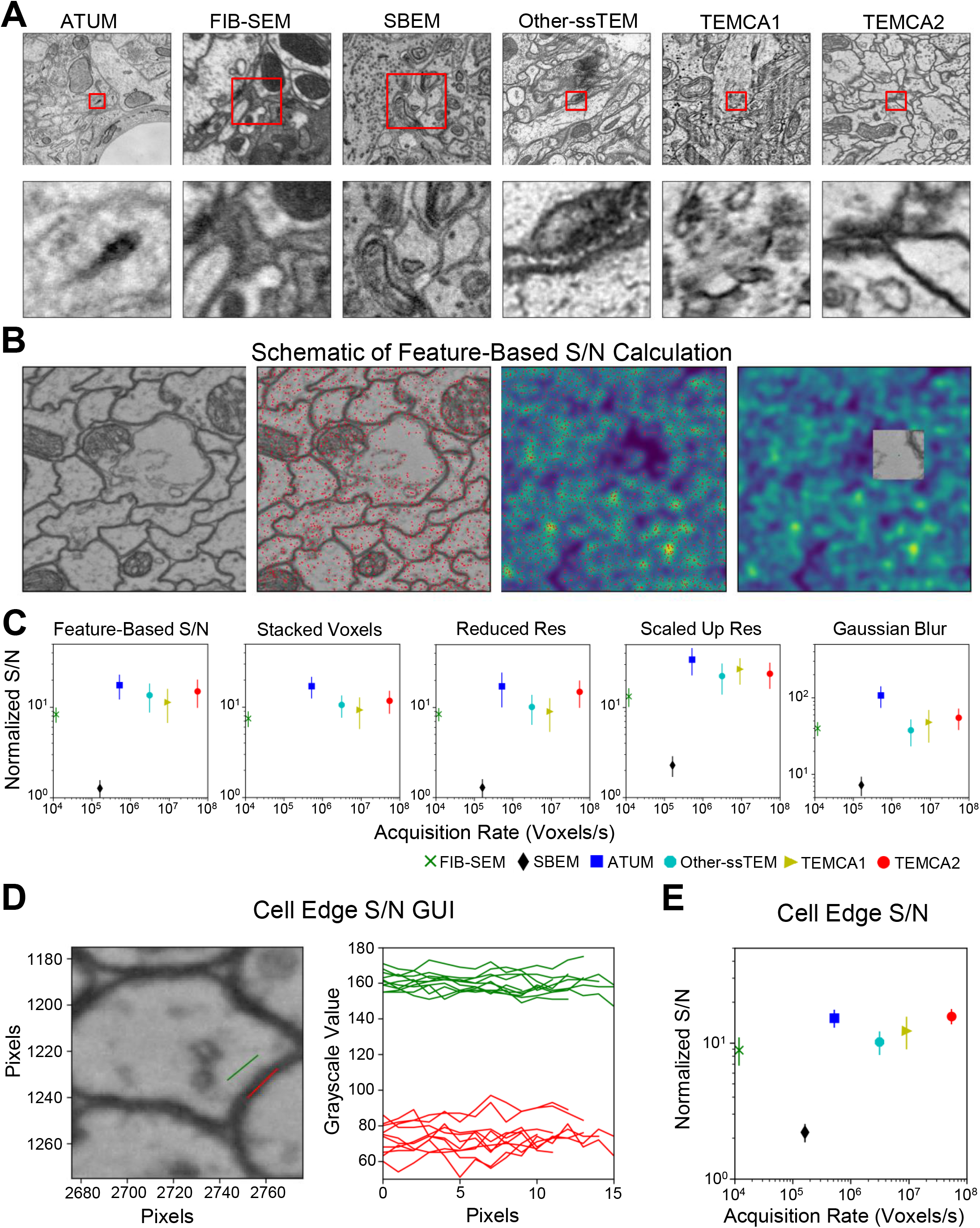
Comparison of S/N Between EM Imaging Methods. Related to Figure 2. (A) Sample images from a variety of EM data sources. From left to right, ATUM-SEM (Kasthuri et al., 2015), FIB-SEM (Takemura et al., 2015), SBEM (Briggman et al., 2011), ssTEM (Takemura et al., 2013), TEMCA1 (Bock et al., 2011), TEMCA2 (this paper). The top row shows images of side length 3 μm while the lower row shows 100 pixel subimages of each. The corresponding areas of these 100 pixel subimages are shown with a black square inside each image. (B) From left to right, a TEMCA2 image sample, the key-points detected in the image, convolution of the key-points illustrating dense and sparse feature regions (purple – low, yellow high), the region of sparse features selected from the TEMCA2 image showing a resin filled area suitable for noise calculation. (C) For all plots points and data sources are as per Figure 2G and Table S3. The normalized S/N versus acquisition rates of a variety of EM techniques (as color coded) are shown for different methods. From left to right, feature-based method as described in (B); Stacked Voxels means that voxels are combined across a layer (SBEM not shown due to alignments not being clear) and across 50 random images; Reduced Res means that voxels correspond to a higher physical size across 100 random images; Scaled Up Res means that voxels correspond to a smaller physical size across 100 random images; Gaussian Blur means that voxels have been blurred with a Gaussian filter across 100 random images. (D) Left, sample image shows a region selected to quantify the noise (green) and a region to quantify the signal (red) for the cell-edge technique. Right, the intensity for noise and signal regions versus number of pixels in each region. (E) Normalized S/N versus acquisition rate as determined via the cell-edge technique across 5 random images from each technique (same color codes as Figure S4C), each of which had 10 regions of background/noise and signal determined. Points and data sources are as per Figure 2G and Table S3.

**Figure S5.**
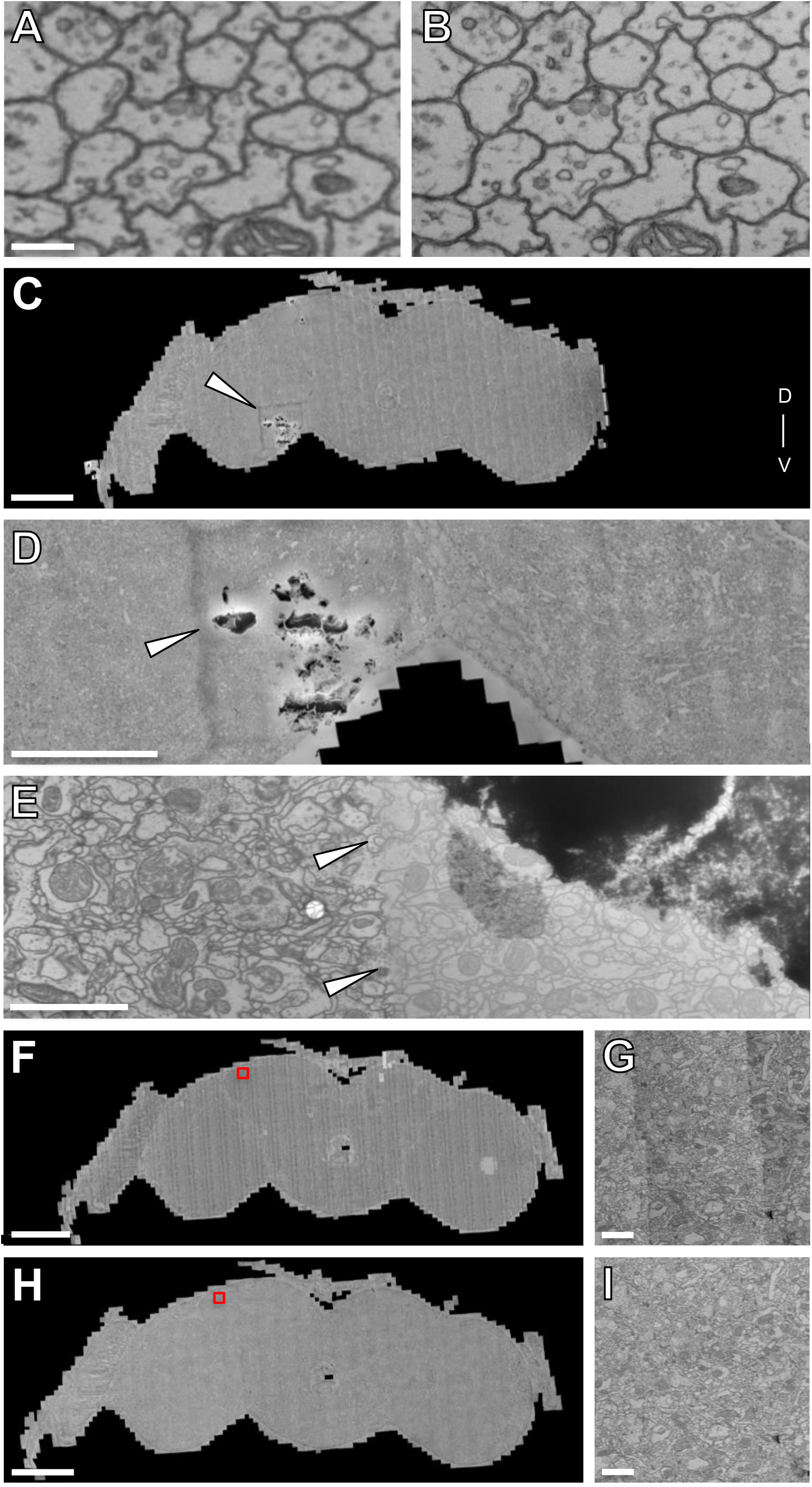
Re-imaging Synapses in Pedunculus, Montaging, and Intensity Correction in 2D. Related to Figure 2. (A-B) Matching ~1.25um wide fields of view in section 3887 from the imaged volume (A) and re-imaged at higher resolution (B). (A) 4nm/pixel; (B) 1.3nm/pixel (C-E) Montaging high-dose and low-dose. Debris present on a section necessitated collection of a small subset of tiles at lower dose than the remainder of the mosaic. (C) The debris and border of the low-dose mosaic can be seen in the context of the entire section. (D) Debris and mosaic boundary are clearly visible. (E) The boundary of the joined high-dose and low-dose mosaics is evident (arrowheads). (F-G) 2D intensity correction. (F) Mosaic of a single section prior to 2D intensity correction. Intensity differences between tiles are evident in (G). (H-I) (H) Same section as in (F) following 2D intensity correction. Intensity differences between tiles are greatly diminished (I). Scale Bars: 200 nm in (A-B), 100 μm in (C), 50 μm in (D), 2 μm in (E), 100 μm in (F, H), 2 μm in (G, I).

**Figure S6.**
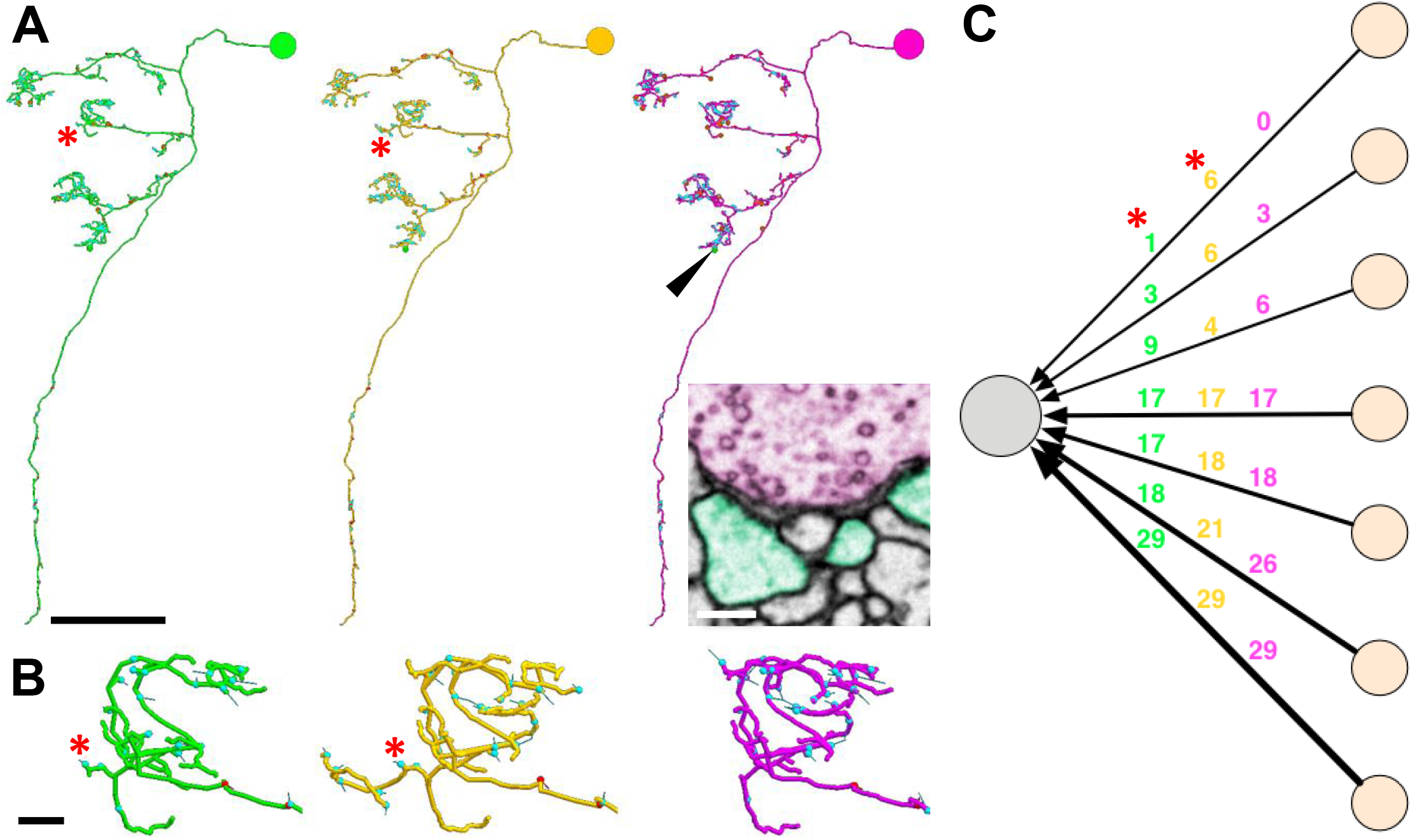
Reliable Tracing. Related to Figure 2. (A-C) Three teams, each comprising 1 tracer and 1 proofreader, reconstructed the same neuron, with each team blinded to the others. (A) Synaptic counts and gross morphologies are comparable across teams. Arrows indicate location of synapse shown IN TEM inset. Asterisks in inset indicate locations of a single KC fingertip postsynaptic to the PN input. (B) Zoom-in to a claw with an input discrepancy across teams. Gold team discovered a *bona fide* process with 6 postsynapses to a putative PN input. Green team discovered 1/5 of the postsynapses on the proximal portion of this process but was not confident to trace the process further. Purple team was not confident to trace this process at all. (C) Network connectivity matched with only 1 inferior input difference shown by red node (PN7). Orange nodes indicate projection neuron bouton inputs to the KC. One putative PN input was missed by team 2, indicated by red box. Colored skeletons (A) and graph nodes (B) and indicate team membership: team 1, green; team 2, gold; team 3, magenta. Scale bars: ~20 μm in (A), 250 nm in (A) inset, ~2 μm in (B).

**Table S1.**
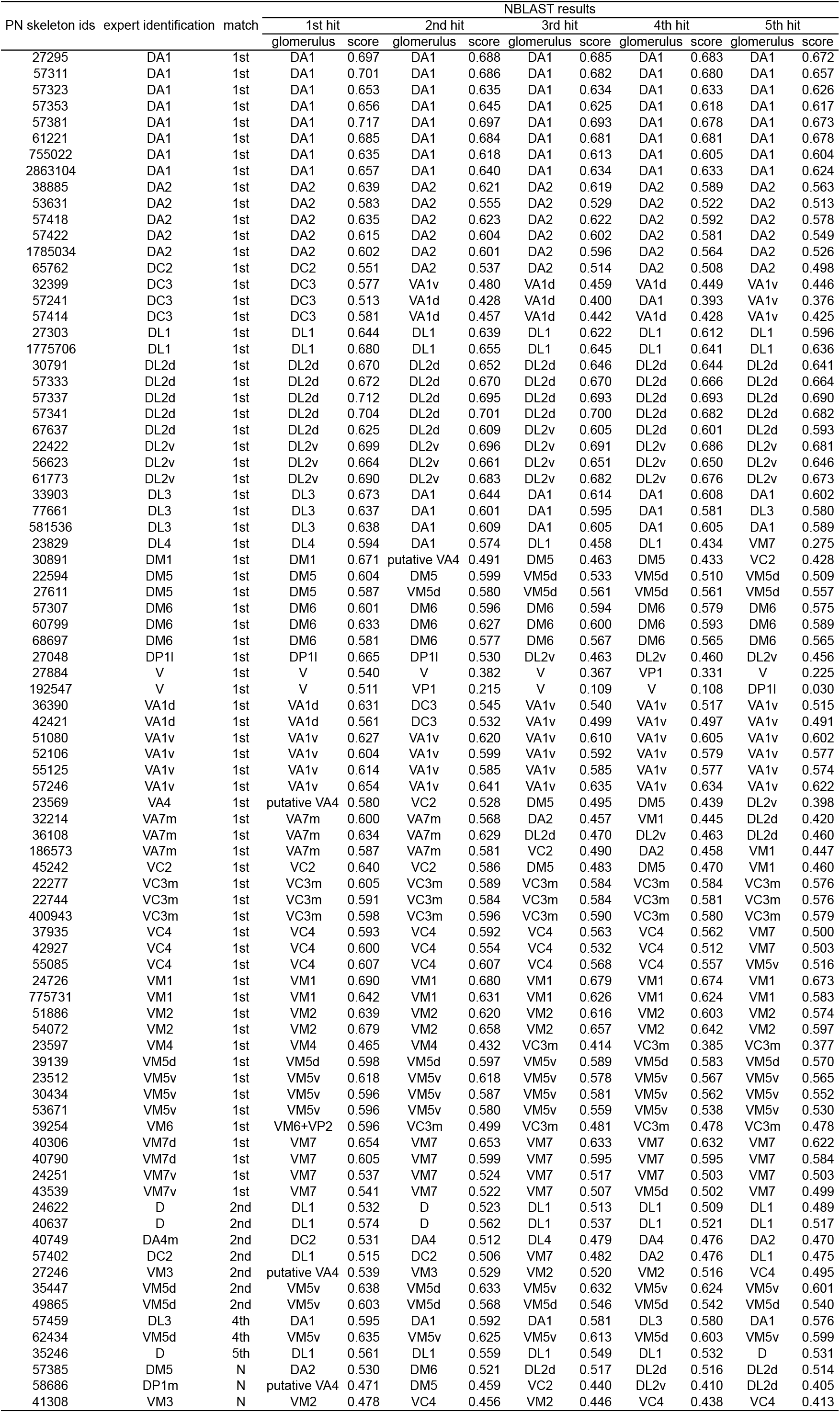
NBLAST Scores for PNs. Related to Figure 4.

**Figure S7.**
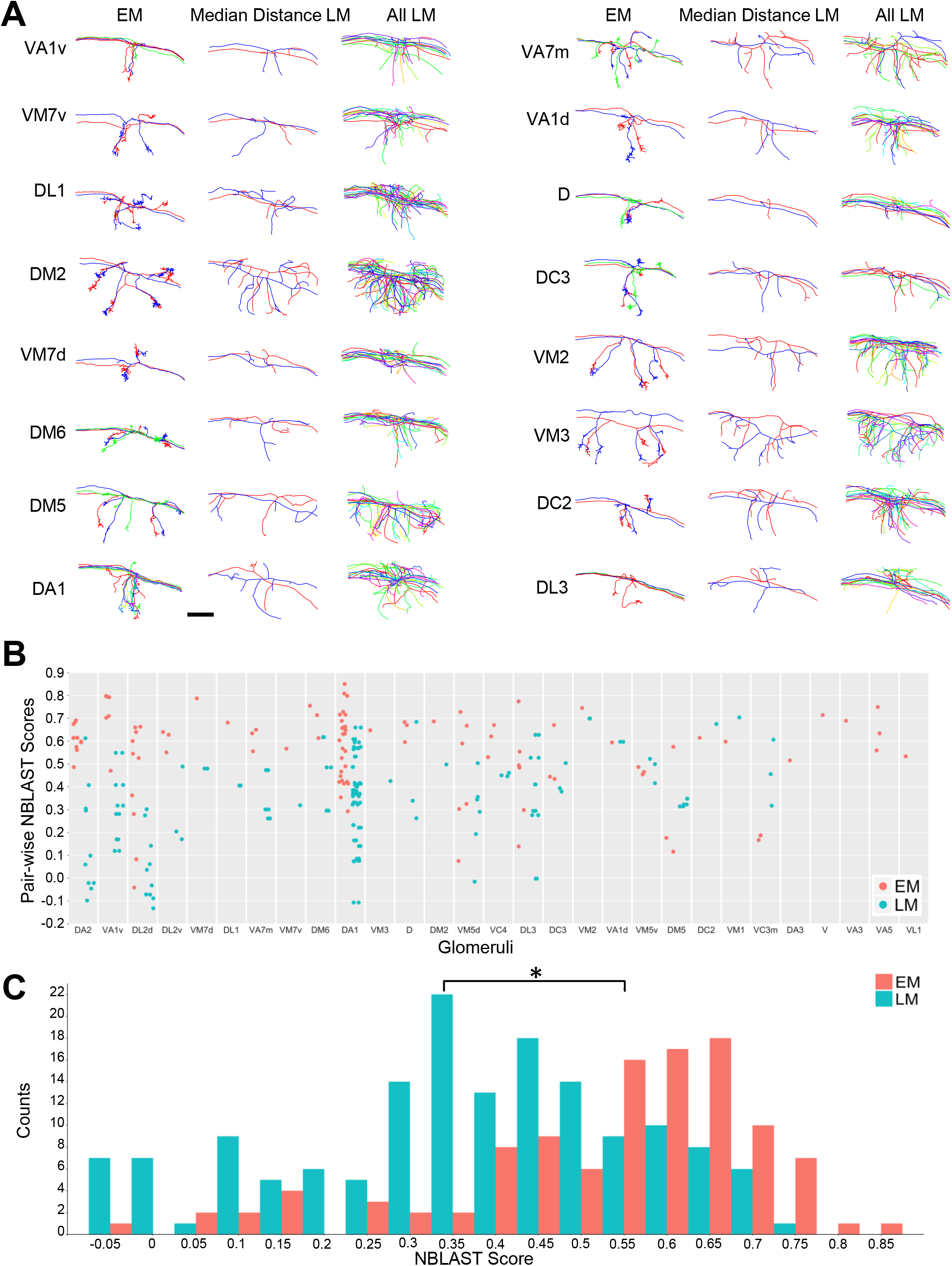
EM versus LM PNs. Related to Figure 5. (A) Qualitative comparison of calyx collaterals of EM versus LM PNs. Only skeletons inside the calyx bounding box is shown, as described in Methods. Only LM PNs from (Jefferis et al. 2007) are used. Glomeruli are ordered by the difference of mean distances between homotypic EM PNs and LM PNs. Left column, EM PNs. Middle column, the pair of LM PNs with median pair-wise mean nearest distances. Right column, LM PNs. (B) Pair-wise NBLAST scores for calyx collaterals of homotypic PNs. Glomeruli are ordered by the difference of NBLAST scores between EM and LM PNs. Each data point represents the NBLAST scores between the calyx collaterals of a pair of PNs from the same glomerulus. (C) Histogram of NBLAST scores across all glomeruli. The average NBLAST scores were significantly higher for PNs in EM than PNs in LM (EM: 0.55 ± 0.19, LM: 0.35 ± 0.19, t test p-value 3.6e-15), indicating that EM PNs are morphologically more similar to each other than LM PNs. Scale bar: ~10 μm in (A).

**Table S2.**
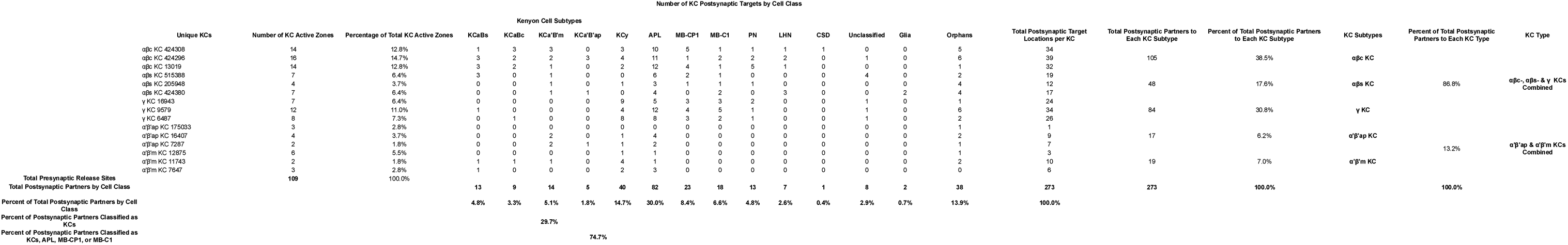
KC Postsynaptic Partners in the MB Main Calyx. Related to Figure 7. Eighty-seven percent of the KC postsynaptic targets are driven by αβc-, αβs-, or γ KCs. The α’β’ KCs drive only 13%. Only 4 neurons are responsible for 75% of the postsynaptic partners. The α’β’ KCs are only presynaptic to other KCs and the APL.

**Movie S4. Whole Brain EM Volume. Related to Figure 1.**

All sections through the dataset are shown. Left, a low-resolution view of the entire section extent. White square is centered on x,y position of the microglomerulus shown in Figure S1A. Right, a zoom-in on a field of view at the center of the white square in the low-resolution view. Section number 5372 shows the microglomerulus of Figure S1A.

**Movie S5. Neuropils Innervated by MB-CP2. Related to Figure 6.**

A previously unidentified neuron, MB-CP2 (orange skeleton), shown inside the whole brain mesh (gray outline) and several other neuropil meshes obtained via LM and registered to the EM volume. MB-CP2 is purely postsynaptic (blue dots) in several regions and both pre- (red dots) and postsynaptic in other regions (first synapse isolation). MB-CP2 innervates the MB (patina), where it is postsynaptic to γ and γd KCs in the MB pedunculus (second synaptic isolation; MB-CP2 skeleton subsequently isolated in blue), and presynaptic to all olfactory KC subtypes at microglomeruli in the MB calyx (second synaptic isolation; MB-CP2 skeleton subsequently isolated in red), where it also receives input from currently unknown cell types. Additionally, it is pre- and postsynaptic in the dAC (compartment mesh not shown), LH, and SLP. MB-CP2 innervates the antlers (cyan), LH (blue), posterior lateral protocerbrum (green), superior clamp (magenta), superior medial protocerebrum (red), superior intermediate protocerebrum (gold), and superior lateral protocerebrum (pink) where it is purely postsynaptic (shown in first synaptic isolation). See Figure 6G for synaptic input/output summary schematic by neuropil. Initial axes: dorsal, top; ventral, bottom. Movie begins from an anterior perspective.

